# The tail of myliobatid rays controls body stability

**DOI:** 10.1101/2025.04.30.651532

**Authors:** Júlia Chaumel, Connor F. White, George V. Lauder

## Abstract

Eagle rays, pelagic eagle rays, cownose rays, and manta rays are the only four batoid families exhibiting oscillatory locomotion, and are characterized by expanded pectoral fins and long, slender tails that can exceed body length. This study investigates whether these tails influence body stability during gliding, when the pectoral fins remain extended and held in position. We first measured relative tail lengths across the four families (*Rhinopteridae, Myliobatidae, Aetobatidae*, and *Mobulidae*). Using two 3D-printed models based on a myliobatid body and a NACA 0012 shape, we tested the effects of different tail lengths (double, equal to disc width, and no tail) on posture and stability across increasing flow speeds (1–4.5 body lengths/second) in a recirculating flow tank. Pitch, roll, and ODBA were recorded via an embedded accelerometer. Our results show that tail length varies among myliobatids, with spotted eagle rays having the longest tails (>4× body length) and giant manta rays the shortest (∼0.9× body length). Models without tails exhibited greater instability, particularly increased roll and ODBA. However, tails longer than body length did not provide additional stability or affect pitch. Although tails offered similar results in both models, the tailless NACA model was more unstable at high speeds than the manta body model. We propose that elongate tails of myliobatids enhance stability by generating hydrodynamic drag and exerting a restoring moment around the center of gravity, thereby damping body oscillations. Tails exceeding 0.9× body length may serve additional functions such as communication, mating, or sensory roles.

**SUMMARY STATEMENT:** What is the function of the long and slender tail in myliobatid rays? We show experimentally that a tail longer than one body length greatly increases hydrodynamic stability during gliding.

## INTRODUCTION

Elasmobranchs are a group of cartilaginous fishes that includes sharks and batoids (rays and skates). Although elasmobranchs comprise only approximately 1500 species, they exhibit a great diversity of body shapes, which are reflective of functional variation in how these groups swim (Carrier et al., 2022; Rosenberger, 2001; Sternes and Shimada, 2020). Sharks typically have fusiform bodies and a heterocercal caudal fin, which oscillates laterally to generate thrust and propel the animal through the water (Smits, 2019; Sternes and Shimada, 2020; Wilga and Lauder, 2004). In contrast, batoids are dorsoventrally flattened with expanded pectoral fins. Although basal batoids (e.g. guitarfishes, electric and shovelnose rays) use their tails for propulsion, most batoids generate thrust using their pectoral fins, enabling the tails to perform a variety of functions that differ across species (Fontanella et al., 2013; Last et al., 2016; Rosenberger, 2001).

Batoid pectoral fin kinematics can be divided into two basic types: undulatory, where waves propagate backward along the pectoral fins, and oscillatory, where the entire pectoral fin moves vertically in unison with a high amplitude (similar to flying birds) (Blevins and Lauder, 2012; Fish et al., 2018; Fish et al., 2016; Fontanella et al., 2013; Heine, 1992; Rosenberger, 2001). Most batoids –including skates and stingrays such as dasyatids – swim by undulatory locomotion, which is correlated with low aspect ratio bodies (diamond or rounder shapes) (Fontanella et al., 2013; Rosenberger, 2001). These batoids are mainly benthic, as undulatory locomotion is considered to provide high degree of maneuverability (Fontanella et al., 2013; Russo et al., 2015). In contrast, oscillatory locomotion is less common, as it is only performed by species from four families of the order Myliobatiformes: *Myliobatidae* (eagle rays), *Aetobatidae* (pelagic eagle rays), *Rhinopteridae* (cownose rays), and *Mobulidae* (manta rays) (Last et al., 2016; White and Naylor, 2016). Species in these families will be referred to here as myliobatids. As oscillatory batoids, myliobatids present triangular body shapes characterized by high aspect ratios (wingspan greater than the length) (Fish et al., 2017; Fontanella et al., 2013; Russo et al., 2015). Oscillatory locomotion with high aspect ratio fins increases locomotor efficiency and is reflected in myliobatid species’ ability for sustained swimming, allowing for long distance migration and extensive vertical movements, features visible in a more pelagic lifestyle than other batoids (Braun et al., 2014; Fish et al., 2017; Fish et al., 2016; Heine, 1992; Klimley et al., 2024; Nosal et al., 2023).

Due to their morphology and corresponding locomotory performance, myliobatids have been a focal point for bioinspired design and extensively studied in fields such as robotics, mechanical engineering, fluid dynamics, and biomechanics (Bianchi et al., 2023; Clark and Smits, 2006; Fish et al., 2016; Liu et al., 2015; Luo et al., 2024; Russo et al., 2015; Xu et al., 2023). These studies have primarily examined how pectoral fins generate thrust and how variations in body morphology influence locomotor efficiency. However, one defining feature of myliobatid rays has been largely overlooked: a slender, whip-like tail with a tapered morphology that can extend up to four times the animal’s body length (Chaumel and Lauder, 2025; Last et al., 2016). This tail is a relatively stiff structure formed by fused vertebrae (the caudal synarcual), which minimizes bending along the tail length, and maintains the tail in a straight, trailing position behind the animal (Chaumel and Lauder, 2025).

The presence of a long, fused vertebral structure within the elongated and tapered tail is unusual among vertebrates. Tapered tails are usually formed by a segmented vertebral column, which allows for a greater range of vertical and horizontal motion (Mallo, 2021). In aquatic locomotion, tapered tails with relatively low vertical surface area are not efficient at generating thrust, as this morphology limits the tail’s ability to displace water (Fish et al., 2021). Instead, tapered tails serve other functions such as providing momentum during swimming, counterbalancing, sensing, or defense (Fish, 1984; Fish et al., 2021; Schwaner et al., 2021; Wake and Dresner, 1967). Similar to other aquatic vertebrates, several batoids also possess long and tapered tails with segmented vertebrae (Last et al., 2016). Although these tails have not been extensively studied, some functions, such as propulsion and defense, can be inferred from associated structures like fins linked to the tails in several species, and a barb in case of the species from the order Myliobatiformes (Hughes et al., 2018; Last et al., 2016; Nishida, 1990). However, the non-segmented, rigid structure, the tapered morphology and the lack of fins on myliobatids ray tails suggest that it may have alternative functions. Recent evidence indicates that the myliobatid tail serves a mechanosensory function, acting like a hydrodynamic antenna, detecting water movements potentially generated by surrounding prey, predators, or conspecifics (Chaumel and Lauder, 2025). However, to date, no studies have explored whether the elongated tail in myliobatids plays any role in influencing locomotion or in providing stability for the moving body.

In this project, we experimentally investigate the hydrodynamic effects of stiff, long and slender tails on myliobatid body stability during gliding with expanded pectoral fins using models. We hypothesize that tails act as passive stabilizers, similar to the stabilizing effect of long, slender tails attached to aerial kites (Borobia-Moreno et al., 2021; Marvin, 1897; NASA, 2021). Our analysis on how tail length influences body stability is confined to gliding behavior when body conformation in rays remains in a relatively static configuration with laterally expanded pectoral fins held in position.

We measured tail and body size across several species of the four oscillatory myliobatid families, to understand the morphological space that these organisms occupy and identify scaling relationships between tail and body morphology. Next, we examined the effects of three different tail lengths on roll, pitch, and overall dynamic body acceleration (ODBA) using accelerometers placed inside a 3D-printed myliobatid body model which was towed in a recirculating flow tank at increasing flow speeds. Finally, as the pectoral fin and body of myliobatid rays are similar in profile to an airfoil (Heine, 1992; Russo et al., 2015), we repeated these experiments with a body model based on a NACA 0012 airfoil cross-section to assess the comparative stability of a foil-shape compared to the body shape of a myliobatid ray. NACA 0012 foils are teardrop-shaped, symmetric foils with no camber, known for their high lift-to-drag ratio and low drag coefficient, and are commonly used in aerodynamic studies and experimental and computational analyses of aquatic propulsion (Ladson, 1988; Lauder, 2015; Lauder et al., 2007; Smits, 2019; Swanson and Langer, 2016; Van Buren et al., 2019; Vandrangi, 2022). Understanding the hydrodynamic role of myliobatid ray tails contributes to understanding the principles that underly body stability during aquatic locomotion and can offer a source of inspiration for the design of robotic devices based on myliobatid rays, in addition to offering new insights into how these species interact with their environment.

## MATERIALS AND METHODS

### Animal morphometrics and density

For morphometric analysis, we measured a total of 76 myliobatids from the four different families: 9 from *Aetobatidae* (*Aetobatus narinari* = 9), 42 from *Myliobatidae* (*Myliobatis californica* = 29; *Myliobatis freminvillei* = 3; *Myliobatis goodei* = 3; *Myliobatis australis* = 2; *Myliobatis aquila* = 1; *Aetomylaeus nichofii* = 2; *Aetomylaeus maculatus* = 1; *Aetomylaeus bovinus* = 1); 8 from *Rhinopteridae* (*Rhinoptera bonasus =* 6; *Rhinoptera brasiliensis* = 2), and 17 from *Mobulidae* (*Mobula mobular* = 5; *Mobula thurstoni* = 6; *Mobula kuhlii* = 4; *Mobula birostris* = 2). The specimens were preserved in 70% ethanol and obtained from several ichthyology collections, including the Harvard Museum of Comparative Zoology (MCZ), Texas A&M University (TCWC), and The Natural History Museum of London (NHM). We measured most mobulids (family *Mobulidae*) from photographs shared by CSIRO’s Australian National Fish Collection in Hobart, Australia. We took dorsal images for each animal, ensuring that the tail was fully extended and clearly visible alongside a scale for accurate measurements and a label to indicate the sex. Embryos and specimens with broken or missing tails were excluded from the study. We analyzed each image in ImageJ software (National Institutes of Health, Bethesda, Maryland, USA) to measure disc width (DW, the distance between the tips of the pectoral fins), body length (BL, the distance from the snout to the base of the tail, between the pelvic fins), tail length (the distance from the base of the tail to the tail tip), and pectoral fin chord (the length of the pectoral fin base) (Fig. 1). Using these measurements, we calculated additional indices such as the tail length to body length ratio, tail length to disc width ratio, estimated body area (DW · chord / 2; which did not include the head, lobes, and pelvic fins) and the aspect ratio (DW^2^ / 2) (Fig. 1).

**Figure 1.**
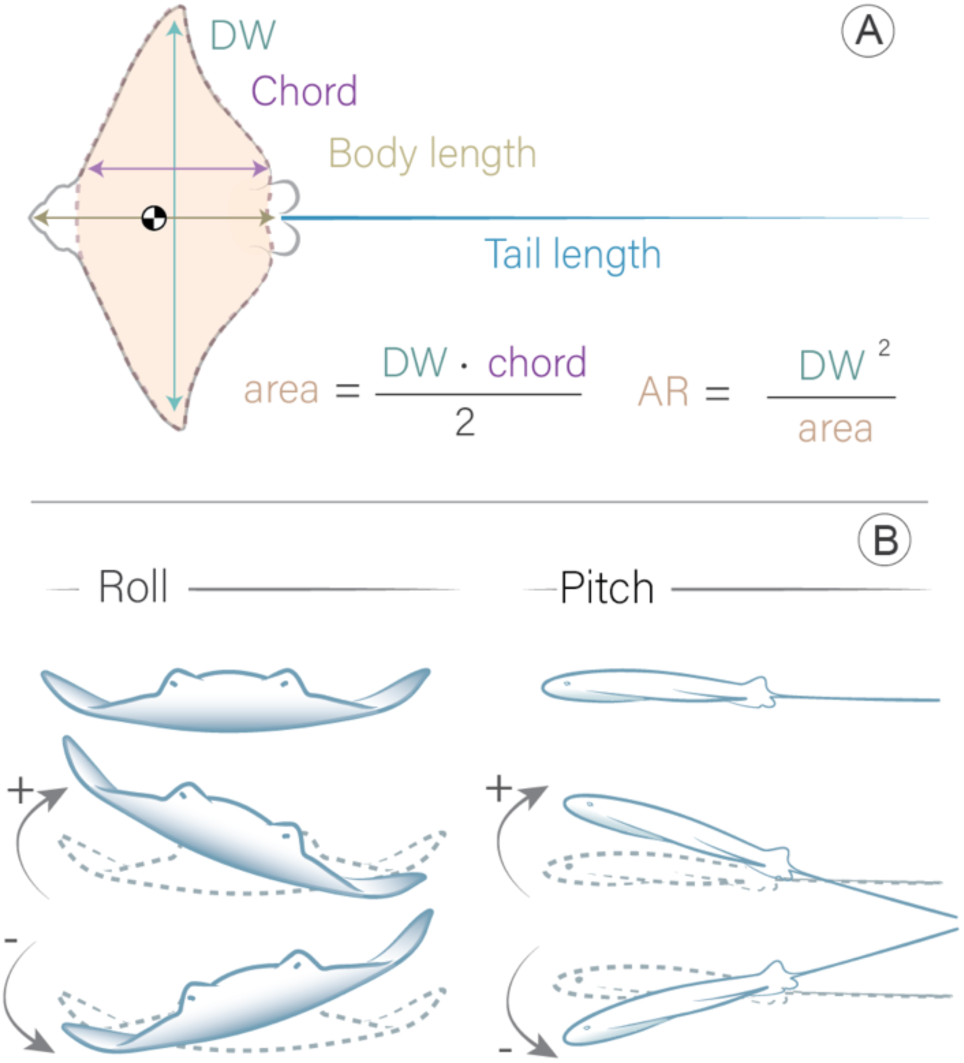
Schematic representation of the measured morphometric variables and ray movement patterns. **(A)** Morphometric variables measured and calculated from specimens, including disc width (DW; distance between the tips of the pectoral fins), chord length (distance between the dorsal and ventral points at the base of a pectoral fin), body length (distance from the head to the base of the tail), and tail length (distance from the base to the tip of the tail). The total body area (excluding the head, lobes, and pelvic fins) was also measured, along with the aspect ratio (AR) of the pectoral fins. The center of mass 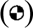 is located anterior to the pectoral fin tips, as described by Fontanella et al. (2013). **(B)** Schematic representation of the myliobatid model’s movements, illustrating positive roll (tilt to the right), negative roll (tilt to the left), positive pitch (upward orientation), and negative pitch (downward orientation).

We calculated the body density of 34 different individuals of a total of 9 myliobatid species from the MCZ ichthyology collection. To determine their density, we measured the weight of the specimens in air (*W_a_*) and in liquid (*W_l_*). To measure the weight in liquid, we suspended the specimens in a bucket filled with 70% ethanol using a balance, ensuring that the specimens did not touch the bottom or sides of the bucket. Based on Archimedes’ principle, the difference between *W*_a_ and *W*_l_ corresponds to the weight of the displaced fluid (buoyant force) (Baldridge Jr, 1970). As specimens were preserved in 70% ethanol (Density = 0.881 g·ml^-3^), the water in their tissues had been replaced by 70% ethanol, resulting in absolute densities less than 1 g·ml^-3^ (the density of water). We assumed that the specimens’ bodies were homogeneous, the volume of the animal did not change during preservation, and the water of their body is fully replaced by ethanol, thus we assumed the weight of the preserved ray in ethanol would be the same as the weight of a non-preserved ray in water. Assuming the density of water to be 1 g·ml^-3^, we calculated the density of the ray using Eqn 1.

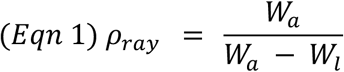

Once the difference between the density of water and density of ethanol are accounted for the estimated densities of individuals ranged from 1.04 to 1.12 g·ml^-3^ and averaged 1.08 ± 0.01 g·ml^-3^ across all measurements.

### Model design and preparation

We designed a myliobatid model using Maya 3D software 2024 (Autodesk Inc., San Francisco, CA, USA) (Fig. 2) that represents a “generalized” myliobatid body morphology. The model was based on 3D models of myliobatids available online (Sketchfab, 2023; Turbosquid, 2023) but altered to ensure left-right symmetry and to incorporate morphological data collected in this study (i.e. observations and measurements of museum specimens). To determine the difference in mass between the body and the tail, we weighed the body and the tail of a *Rhinoptera bonasus* specimen separately (specimen MCZ_49097 from the Harvard Museum of Comparative Zoology). For this specimen, the tail accounted for 1.3% of the total mass of the animal (body mass = 1543g, tail mass = 21g), and in our experiments, a tail of comparable relative length represented 2.3% the total mass (body mass = 47.3g, tail mass= 1.1g). The myliobatid body model was designed with a disc width of 15 cm and a body length of 10 cm (Fig. 2A, C).

**Figure 2.**
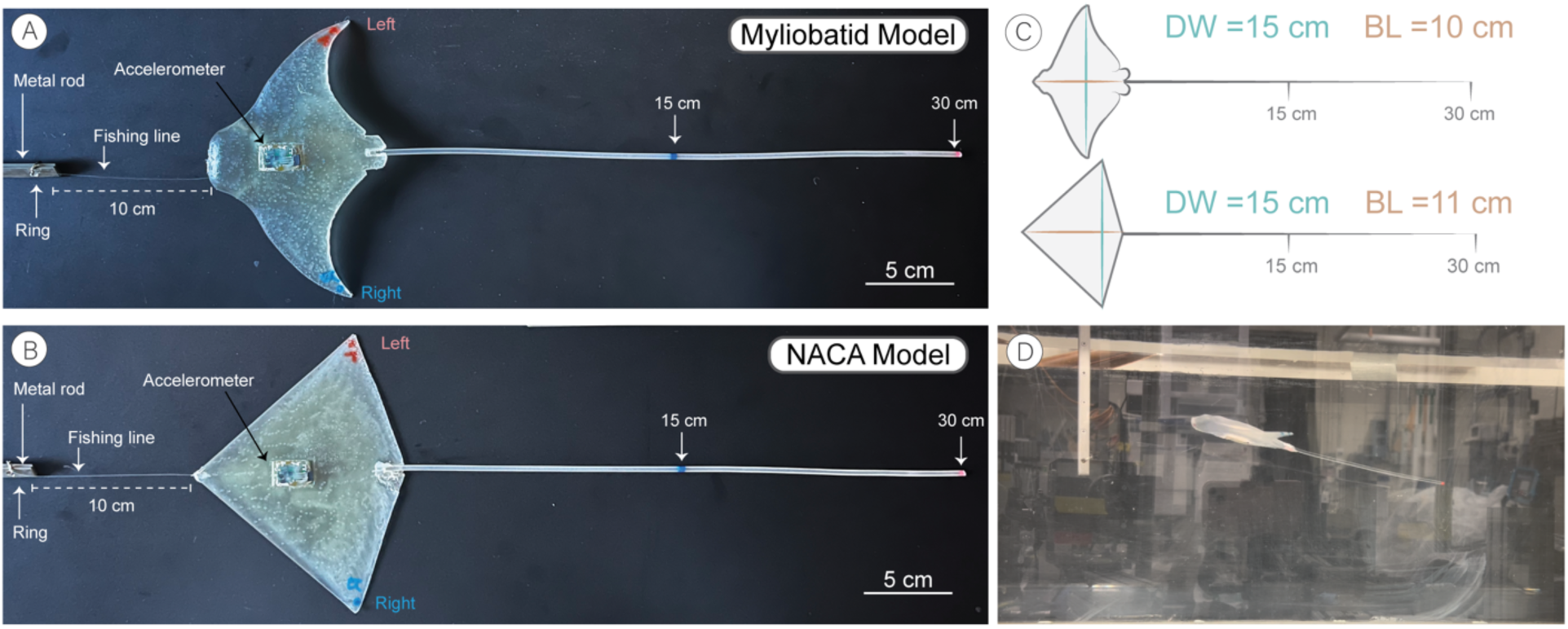
Models and experimental setup. **(A)** Myliobatid model used in the experiments, with a tail marked at 30 cm (2:1 disc width) and at 15 cm (1:1 disc width). (**B)** NACA 0012 model with the tail marked at 30 cm (2:1 disc width) and at 15 cm (1:1 disc width). Both models were attached to a metal rod using an 8-pound fishing line of 10 cm attached to a ring and tied to the hole visible at the bottom of the vertical rod. The accelerometer was placed on the ventral side of the models. **(C)** Schematic representation of both models (disc width, DW, and body length, BL) and their respective tails. **(D)** Flow tank experimental setup showing the myliobatid model with a 15 cm tail suspended in the recirculating flume with flow. The model has assumed a positive angle of attack and positive pitch at this speed.

Additionally, we generated an abstracted myliobatid model (NACA model) with a simplified and known geometry using Maya 3D software 2024. The ‘NACA model’ employed a NACA 0012 (National Advisory Committee for Aeronautics) cross-sectional profile in the streamwise dimension. We selected the NACA 0012 profile because it is symmetrical, maintains a high lift to drag ratio over a range of angles of attack, and has been previously used as a biomimetic shape to understand aquatic locomotion (Aasha et al.; Ladson, 1988; Mei et al., 2021; Prabu et al., 2024). To generate a delta-like shape (similar to a myliobatid-like body shape), the cord length (and thus thickness) of the model decreased towards the wing tips, and the location of maximum wingspan matched that of the Myliobatid model at ∼77% of the body length. The NACA model was similar in weight (body mass = 43.7g), where the tail represented 2.5% of the total mass, disc with a 15 cm DW and a body length of 11 cm (Fig. 2B, C).

We designed both 3D-printed models with several internal cavities essential to experimental testing. On the ventral surface at the approximate center of mass (which correlates with the position of the center of mass measured in myliobatids, (Fontanella et al., 2013)), we placed a rectangular cavity (2.3×1.3x 0.9 cm) in the model to house an accelerometer data logger. We added a small cavity (0.7 cm diameter) at the rostrum’s (anterior) midpoint for attaching a tether to secure the model during flow tank experiments. Finally, we created a small (0.3 cm diameter) cylindrical cavity along the posterior edge of the model between the pelvic fins to attach tails of varying length (Fig. 2A,B).

We 3D-printed the models with the stereolithography printer FormLabs 3+ (FormLabs, Somerville, MA, USA). The resin used was *Elastic50A* V2, because its density (1.01 g / ml^3^) closely resembles that of measured myliobatid body densities (calculated above). Once printed, we inserted an accelerometer data logger (Axy-5, TechnoSmart, Rome, Italy) into the ventral cavity of the model and fixed in place using hot glue (Fig. 2A, B). Prior to each trial, we attached a new tail of 30 cm in length to the cavity between the pelvic fins and fixed it in place using hot glue. We made the tails using a semitransparent, flexible thermoplastic polyurethane filament NinjaFlex (NinjaTek Inc., Manheim, PA, USA) with a diameter of 0.3 cm. We straightened the tails using heat with a heat gun to remove any residual curvature. We placed a blue mark at the 15 cm midpoint to mark half tail length, and a red mark at 30 cm to indicate the tail end (Fig. 2A, B). Finally, we secured a tether (10 cm-long, Sufix 8-pound monofilament line; Greensboro, NC, USA) to the rostral cavity of the model using hot glue. We attached the other end of the tether to a ring mounted on a rigid metal rod used for securing the model in the flow tank (Fig. 2 A,B,D). The ring provided the model with a greater range of motion.

### Flow tank testing

We tested models with and without tails in a recirculating flow tank with varying flow speeds and tail lengths. The flow tank had a 28×28×66cm (width, height, length) cross section and has been extensively used in previous work (Fig. 2D) (Di Santo et al., 2017; Lauder and Tytell, 2005). A metal rod was vertically fixed in place so that the tether attachment was ∼18cm from the bottom of the tank. This allowed the models to rest on the bottom of the tank when there was no flow.

During the first set of experiments, we exposed both models (myliobatid and NACA) with a 30cm tail (2x DW) to a flow speed of 7.5 cm·s^-1^ (0.75 BL·s^-1^) which was increased by 0.98 cm·s^-1^ (or every 25 rpm for the propeller) every two minutes in a stair step protocol, concluding at 46.8 cm·s^-1^ (4.6 BL·s^-1^). We set the minimum speed at 7.5 cm·s^-1^ because it was the lowest speed at which a model generated sufficient lift so that it was no longer resting on the bottom. After the maximum speed was reached, we cut the tail to 15 cm (1x DW) and we repeated the speed sequence. Finally, we trimmed the tail to 0 cm (no tail) and repeated the experiments with the same speed sequence. We conducted this experimental protocol four times for each model (NACA and the myliobatid). We recorded video data for all trials with a lateral-view camera.

To assess how small variations in tail length affected model stability, we conducted a second set of experiments. We placed the models in the flow tank at a constant speed of 14.1 cm·s^-1^ (1.4 BL·s^-1^). We chose this speed as all models generated sufficient lift to raise the model to the mid-water level (Fig. 2D) and because of its biological relevance, falling within the known swimming speed range of myliobatid species (Fish et al., 2016). During the experiment, we gradually reduced the model’s tail length from 30 cm to 0 cm by cutting 1.5 cm from the tail every 2 minutes (e.g. 30, 28.5, 27, 25.5 etc.). We repeated this experiment three times for each model. Additionally, we later took high-speed camera footage from both lateral and bottom views of models in the flow tank at a speed of 1.4 BL·s^-1^ to allow subsequent analysis of body motion.

### Data analysis

During all experiments, the embedded acceleration data logger recorded pressure and temperature at 1Hz, and tri-axial acceleration at 100Hz. At the conclusion of each experimental trial, we downloaded the datalogger data. We aligned the dataloggers within the models so that the accelerometers x-axis corresponded to the rostral-caudal axis, the y-axis corresponded to the medial-lateral axis, and the z-axis corresponded to the dorsal-ventral axis of the models. The data logger records proper acceleration, or the acceleration relative to free-fall, which can be decomposed into two components, static and dynamic acceleration. Static acceleration is acceleration due to gravity (9.8 m·s^-2^) and as this vector is fixed in the global frame it can be used to reconstruct the posture (pitch and roll) of the model. Body pitch is the angle between the horizontal and the model’s rostral-caudal axis (i.e. rotation around y-axis), while body roll is the angle between the model’s medio-lateral axis and the horizon (i.e. rotation around the x-axis). We calculated pitch and roll of the models at each moment in time using Eqn 2 and 3

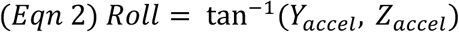

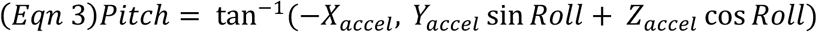

A positive pitch angle indicates that the front of the model is tilted upwards, while a negative pitch angle indicates that the model is tilted downward (Fig. 1B). A positive roll value corresponds with a model tilting to the left, while a negative roll value corresponds to the value tilting to the right (Fig. 1B). However, the recorded acceleration values additionally contain dynamic acceleration, or the acceleration due to changes in speed and orientation of the model. We calculated static acceleration using a 3 second running mean (300 point) of the raw acceleration data. We calculated dynamic acceleration as the remainder when subtracting the static acceleration from the raw acceleration (Shepard et al., 2008). From dynamic acceleration we calculated overall dynamic body acceleration (ODBA, Eqn 4), a metric that captures the degree of three-dimensional movement of an object and has been extensively used to capture animal activity levels (Gleiss et al., 2011).

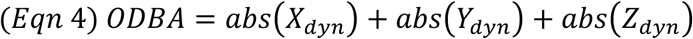

For statistical analysis, we collapsed the tag data into two-minute bins based on speed and tail length for each trial. However, we removed the first 20 seconds of each treatment after manipulation (i.e. speed change or tail cutting) so that only the steady state behavior was captured. From each trial bin (speed, tail and model type), we summarized the mean body pitch and body roll to understand the orientation and position of the model. To quantify the stability of the model, we calculated several metrics including the mean ODBA, as well as range (maximum – minimum) of roll and pitch values experienced.

### Statistical analysis

To examine morphological variations across myliobatid species, we analyzed a series of linear models, and to examine the relationship between tail length and body size we conducted regressions of tail length against body length. Species with less than three individuals were not included in the statistical analysis, except for *M. birostris.* In these models, we incorporated an interaction between body size (continuous variable) and species (discrete factor), allowing each species to have their own slope and intercept. To test if the tail length varied with the body shape among myliobatid species, we analyzed the standardized tail length (tail length / body length) against the aspect ratio (Fig. 1) for each myliobatid species with ANOVA. To determine which species contained significantly different standardized tail length relative to body shapes (standard tail length, aspect ratio), we conducted post hoc comparisons using the Tukey HSD test.

To analyze how flow speed and tail length impacted body position (pitch, roll) and stability (ODBA, pitch range, roll range) metrics, we constructed linear models. Body position and stability metrics were response variables, while tail length, flow speed, and the interaction between tail length and flow speed were the predictor (independent) variables. Tail length and flow speed were classified as discrete factors and not continuous variables. Post hoc analysis included pairwise comparisons (t-test) using planned contrasts (n=24), so that only comparisons between tail lengths at the same speed were included. We adjusted the significance value of post hoc tests to account for multiple testing using the Bonferroni correction. We constructed the statistical models separately for the myliobatid and NACA models.

To determine if there were significant differences between the NACA and myliobatid models, we constructed a separate series of statistical models. First, we conducted linear models with positional and stability metrics as response variables, while the predictor (independent) variables were model type (myliobatid, NACA), flow speed, and the interaction between them. Then, we conducted separate models for each tail length (30, 15, and 0cm). We used t-tests as post hoc pairwise comparisons so that only models were compared with the same tail lengths and at the same speed.

Models with 30 and 15 cm tails had their tail in contact with the bottom of the tank at the lowest tested speed. As this altered their movement, all 7.5 cm s^-1^ speed measurements were excluded from analysis. In order to facilitate comparison across studies, we have presented speeds in units of body lengths per sec BL·s^-1^, by dividing the flow speed by the model’s body length. All data and statistical analyses were conducted in R (R Foundation for Statistical Computing, Vienna, Austria, Version 4.4.2).

## RESULTS

### Myliobatid morphology

We analyzed how tail length varied with body length, disc width, and aspect ratio across 15 different species (n = 76 individuals) from the four oscillatory families of myliobatiforms (*Mobulidae*, *Rhinopteridae*, *Myliobatidae*, and *Aetobatidae*). The specimens available were mainly juveniles and presented a large variation in tail length (22–162 cm), body length (9–136 cm) and disc width (16–338 cm, Fig. 3). Mobulids were the largest myliobatids, with *Mobula mobular* (devil ray) possessing the longest tail (TL = 162 cm) and *Mobula birostris* (giant manta ray) the largest body size (BL = 136 cm, DW = 338) (Fig. 3). Across all included species, tail length increased with body length (F_1,49_ = 671, p < 0.001) and disc width (F_1,49_ = 558, p < 0.001). However, the rate at what the tail increased varied among myliobatid species (BL: F_8,49_ = 4.3, p < 0.001; DW: F_8,49_ = 5.5, p < 0.001) (Fig. 3D).

**Figure 3.**
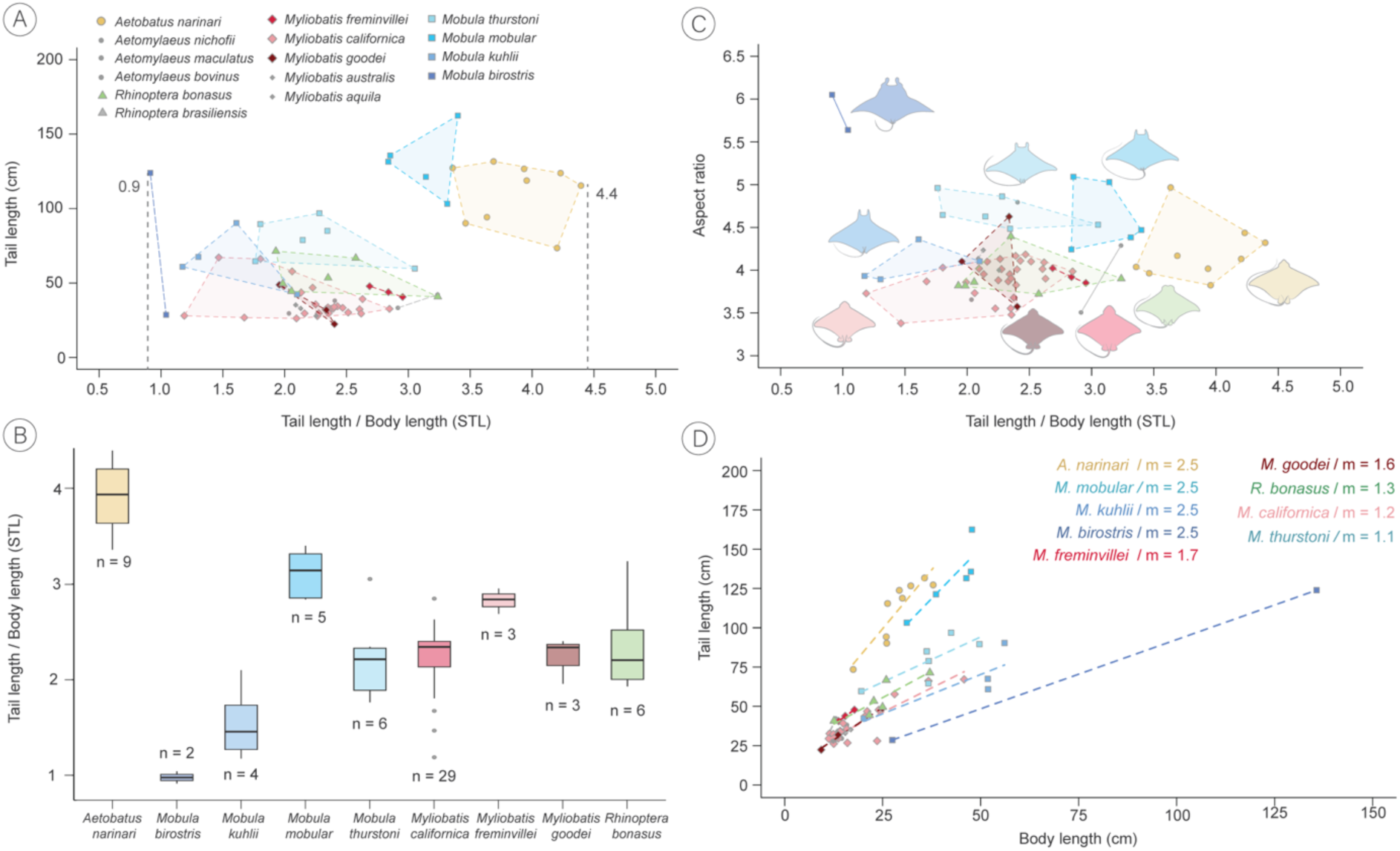
Tail length variation across myliobatid species. **(A,B)** Variation in standardized tail length (STL; tail length/body length) among myliobatid species (n = 76). Only species with more than two individuals were statistically analyzed, except for *Mobula birostris* (n = 2). Statistically analyzed species are highlighted in different colors, while non-analyzed species are shown in grey. STL varied across different myliobatid species (ANOVA, p < 0.001), with *M. birostris* having the shortest STL (0.9 BL) and *Aetobatus narinari* the longest (4.4 BL). *Mobula thurstoni* exhibited the longest absolute tail length recorded. **(C)** Relationship between STL and aspect ratio (AR) across myliobatid species. Mobulids consistently showed the highest AR compared to other myliobatids. *M. birostris* had the highest AR and *Myliobatis californica* had the lowest AR. In contrast, *A. narinari* displayed a low AR but the longest tail. **(D)** Correlation between tail length and body length for species with more than two specimens measured. For all species, tail length increased with body length (ANOVA, p < 0.001) but at different rates among species (ANOVA, p < 0.001). *A. narinari*, followed by *M. mobular*, the tail length increased at higher rate relative to body length (higher *m* values), contrasting with *M. birostris*, which tail length increased at the lowest rate relative to body length.

To calculate the standardized tail length (STL) across myliobatid species, we calculated the tail-to-body length ratio (tail length / body length). Our results showed that the STL significantly differed among species (ANOVA, F_8,58_ = 28, p < 0.001), ranging from 0.9 in *M. birostris* (meaning that the tail is 0.9 times the body length) to 4.4 in *Aetobatus narinari* (spotted eagle ray) (where the tail is 4.2 times the body length) (Fig. 3A,B). In fact, *Aetobatus narinari* had significantly longer tails (STL = 3.8 ± 0.3 BL) than any other species (Tukey HSD, p<0.01). While *Mobula birostris* (giant manta) only had two observations precluding robust statistical analysis, these were the two shortest tails observed in this study (STL = 0.9 ± 0.1 BL) (Fig. 3A,B). For species with more than five observations, the range between the shortest and longest tail within a species always exceeded 1 body length (Fig. 3). Regarding the STL for other species, myliobatids from the families Rhinopteridae *(Rhinoptera bonasus*_STL_ *=* 2.4 ± 0.5 BL), Myliobatidae (*Myliobatis californica*_STL_ = 2.2 ±0.4 BL), and Mobulidae (*Mobula thurstoni*_STL_ = 2.2 ± 0.5 BL) displayed overlapping tail lengths that did not differ from each other (Tukey HSD, p > 0.9). Notably there was variation within mobulid, with *M. mobular* (STL = 3.1 ± 0.3 BL) having significantly (Tukey HSD, p = 0.003) longer tails than *M. thurstoni* (STL = 2.2 ± 0.5) and *M. kuhlii* (STL = 1.5 ± 0.4). With the exception of *M. birostris*, species with less than three individuals were not statistically analyzed but plotted in the graphs (grey points in Fig. 3A,C).

Similar to STL, there was significant variation in aspect ratio (AR) across species (ANOVA, F_8,58_ = 19, p<0.001) (Fig. 3C). *Mobula birostris* had the highest aspect ratio (5.8 ± 0.3) and two other species of *Mobula (M. thurstoni*_AR_ = 4.7 ± 0.2; *M. mobular*_AR_ = 4.6 ± 0.4) presented a higher AR than other myliobatids (Tukey HSD, p < 0.1). However, the aspect ratio of *M. kuhlii* (AR = 4.1 ± 0.2), *A. narinari* (AR = 4.2 ± 0.3), *R. bonasu*s (AR = 3.9 ±0.2), and *Myliobatis* sp. (*M. freminvillei*_AR_ = 3.9 ± 0.1; *M. californica*_AR_ = 3.9 ± 0.2), did not differ from each other (Tukey HSD, p > 0.9) (Fig. 3C).

By measuring the weight in air and weight in ethanol we were able to estimate the density of 10 myliobatid species (n=34 individuals). There was no significant difference in density among species (F_10,23_ = 0.9, p = 0.51) or with body length (F_1,28_ = 2.5, p = 0.11). However, we were only able to measure individuals < 33.7 cm DW. These measurements provided an approximate reference for the models tested in the flow tank.

### Model posture

At the lowest speed (10 cm·s^-1^ or 1 BL ·s^-1^), both the NACA and myliobatid models displayed their maximum pitch value (tilted up posture, with a positive body angle of attack to oncoming flow) (Fig. 4–6, Table 1; Table S1). As flow speed increased, the pitch of the models significantly decreased (myliobatid: F_15,134_ = 78, p<0.01, NACA: F_15,132_ = 99, p < 0.01) (Fig. 4–6, Tale 1; Table S1) and models were approximately horizontal (pitch < 5°) once speeds reached 3 BL ·s^-1^ (30.5 cm·s^-1^) (Fig. 4–6, Table 1; Table S1). While the NACA models roll (roll < 3°; Table S1) did not change with flow speed (F_15,132_ = 0.52, p = 0.92) (Fig. 5), once speed exceeded 3.0 BL·s^-1^ the myliobatid model displayed a progressive roll to the right-hand side (F_15, 132_ = 18.6, p <0.001) (Fig. 4; Table 1). As the speed increased, the Myliobatid model’s roll could exceed 45° and it would turn upside down (roll=180°) at speeds faster than 5.7 BL·s^-1^ (57.7 cm·s^-1^) (Fig. 4). As the myliobatid model’s roll increased, the estimated pitch of the model also displayed minor increases (pitch < 5 °) (Fig. 4, 7b, Table 1). Due to this rolling behavior, speeds higher than 4.7 BL ·s^-1^ (47 cm·s^-1^) were not included in the statistical tests for any of the models.

**Figure 4.**
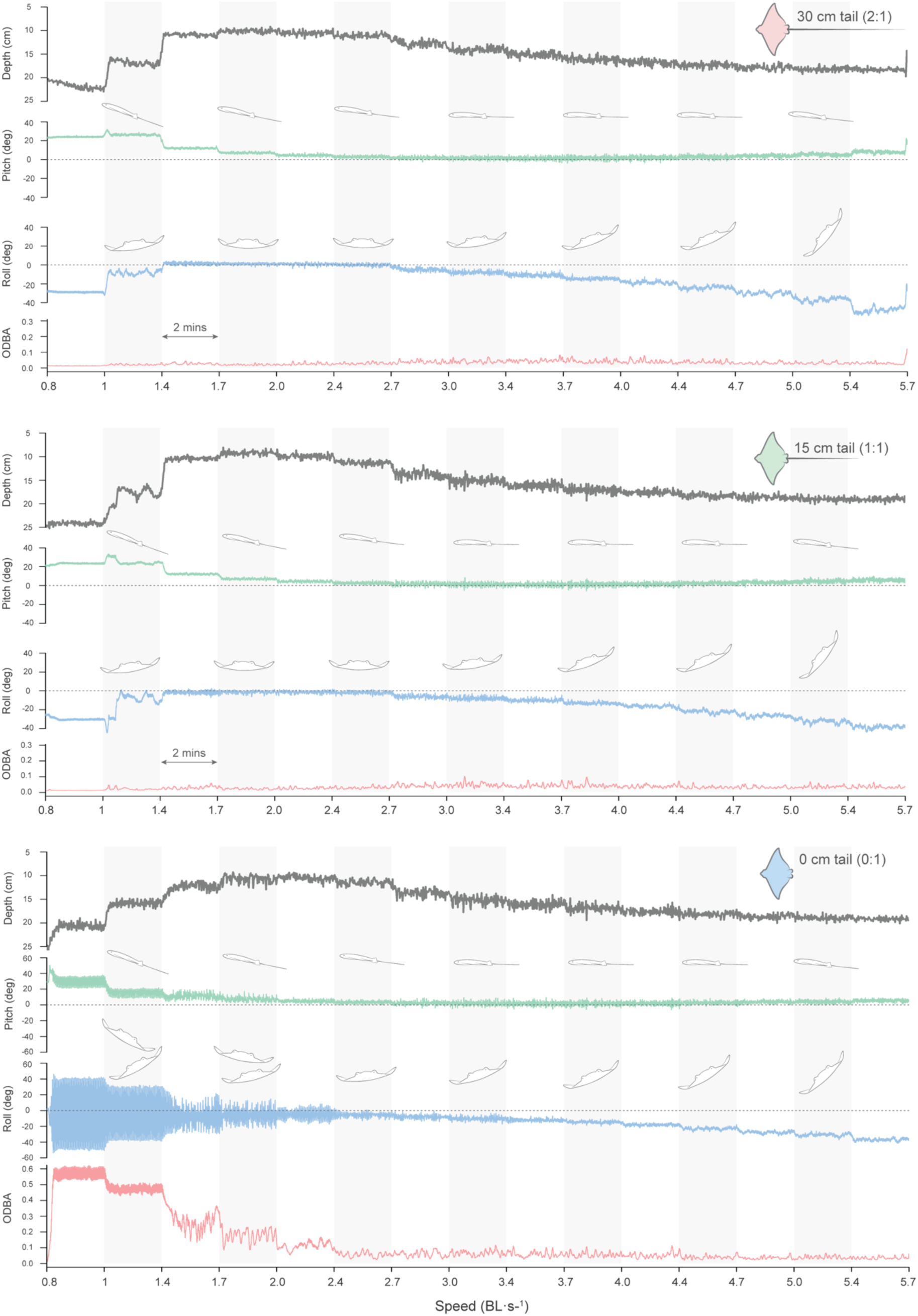
Depth, pitch, roll, and ODBA variation of the myliobatid model across increasing flow speed and different tail lengths in a single trial. Horizontal bars (grey and white) indicate flow speed intervals, each maintained for 2 minutes. The myliobatid model without a tail exhibited a significant increase in pitch, roll, and ODBA. Notably, the model tended to tilt to the right as flow speed increased.

**Figure 5.**
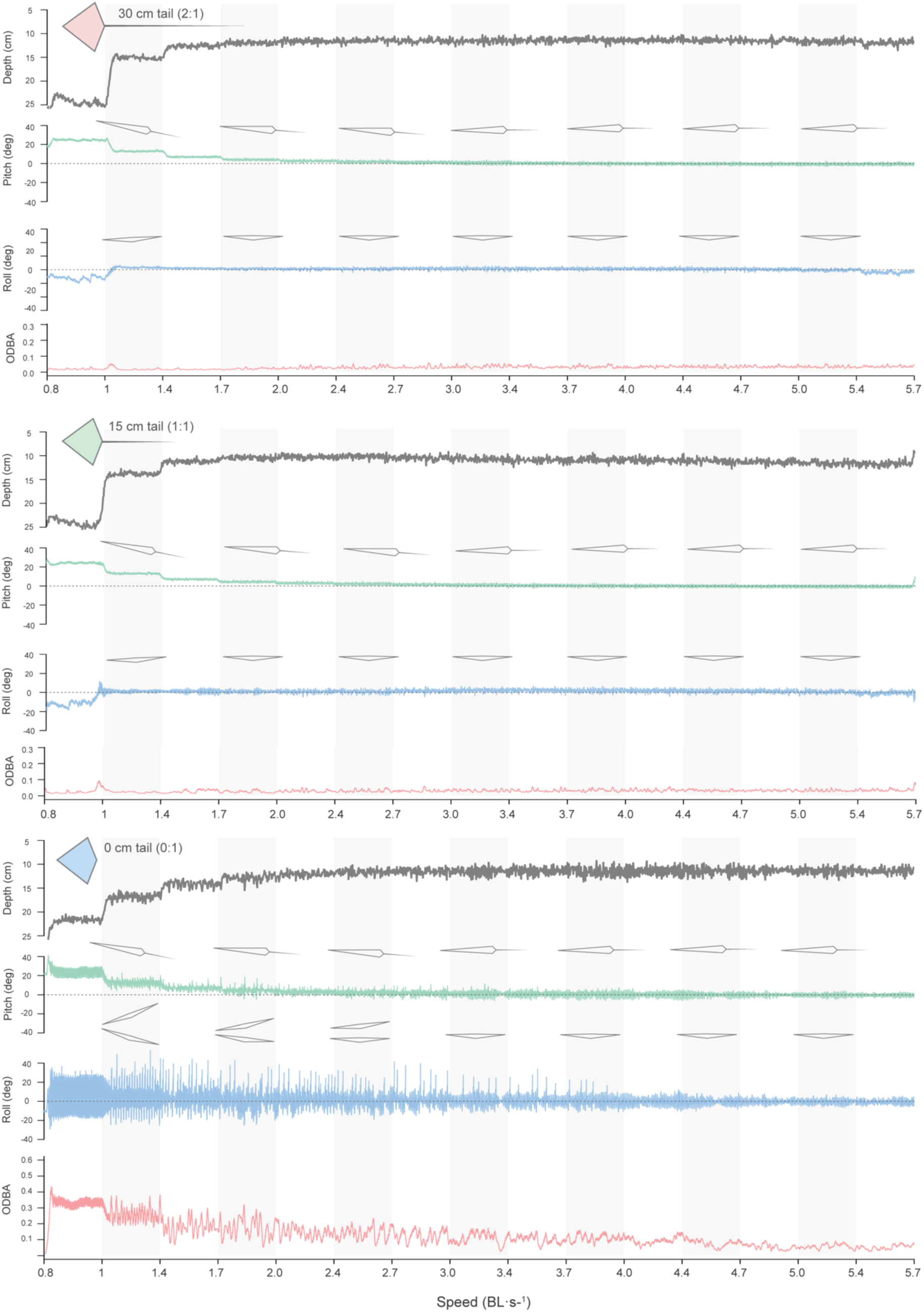
Depth, pitch, roll, and ODBA variation of the NACA 0012 model across increasing flow speeds and different tail lengths in a single trial. Horizontal bars (grey and white) indicate flow speed intervals, each maintained for 2 minutes. The NACA model without tail had a higher pitch, roll, and ODBA that decreased with higher flow speeds than the myliobatid model. Note that this model does not tilt to one side as the speed increases.

**Figure 6.**
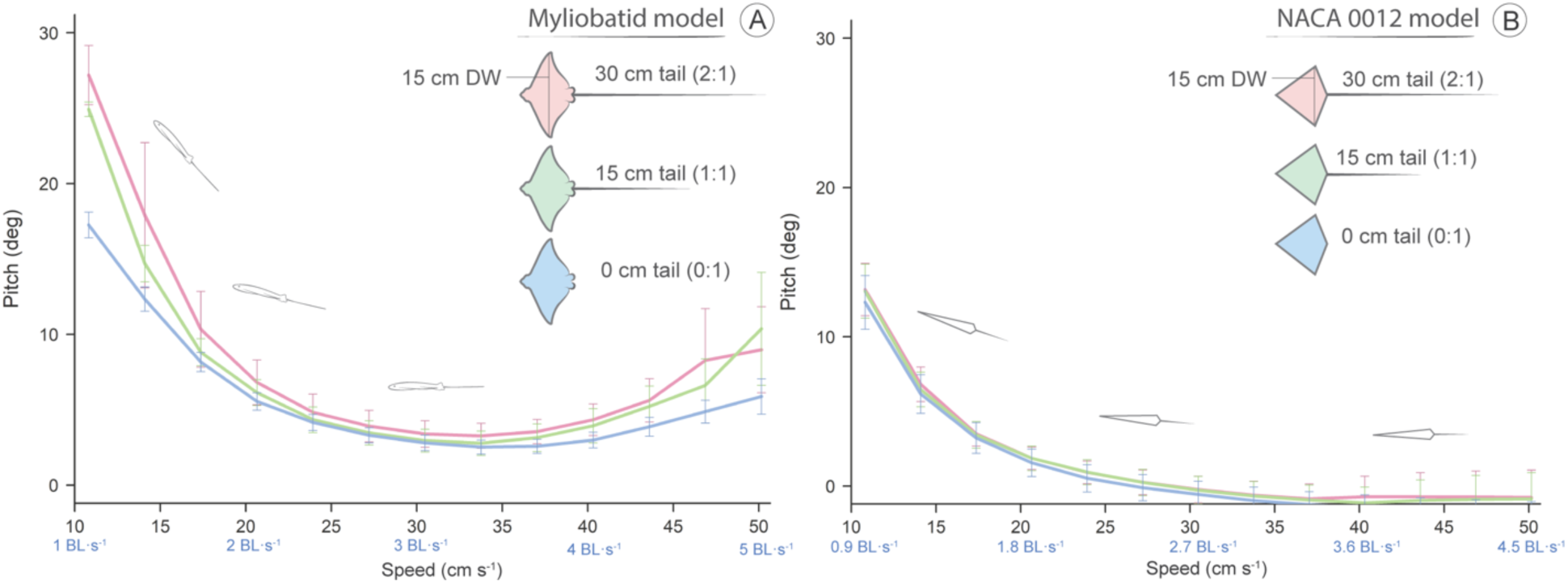
Variation of mean pitch values across flow speeds and tail lengths. Lines of different colors on the graph represent each tail length (30, 15, 0 cm). Error bars indicate the standard error based on four repeated trials for each tail length. Flow speeds are indicated as cm·s^-1^ and BL· s^-1^ (body length). Overall, both models presented their maximum pitch value at lower flow speeds and, as the flow speed increased, the pitch significantly decreased (ANOVA, p < 0.001). **(A)** For the myliobatid model, both tail lengths (30 and 15 cm) produced higher mean pitch values than the model without tail (Tukey HSD, 0:15cm: p < 0.001; 0:30cm: p = 0.005). However, as flow speed increased, no significant differences were observed between different tail lengths (Tukey HSD, p > 0.1). **(B)** The NACA 0012 model presented no differences in pitch average among all tail lengths (ANOVA, p = 0.38) and displayed a lower pitch angle than the myliobatid model (ANOVA, p < 0.01).

**Figure 7.**
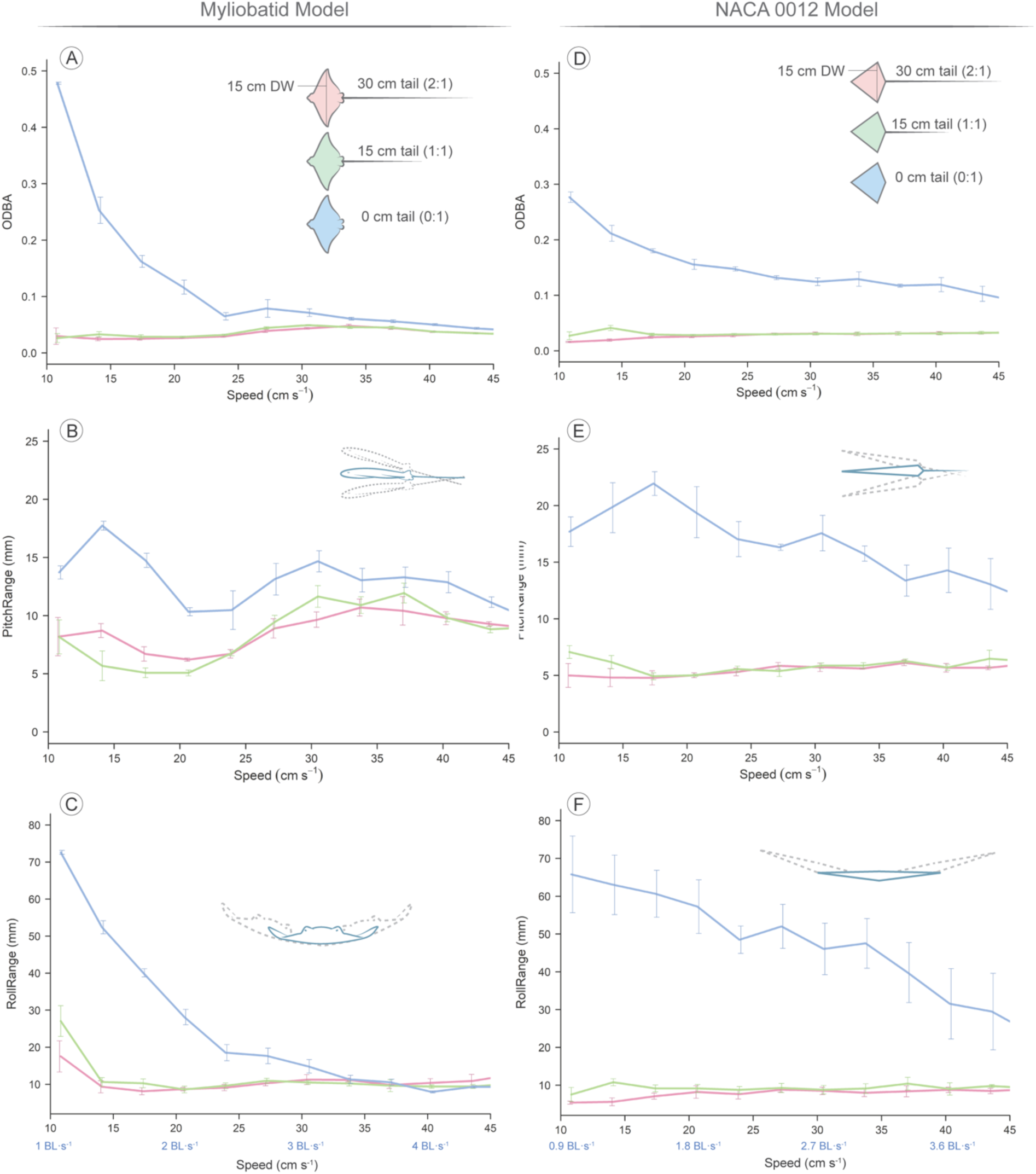
Variation in pitch range, roll range, and ODBA for both models across tail lengths and flow speeds. Lines of different colors on the graph represent each tail length (30, 15, 0 cm). Error bars indicate the standard error based on four repeated trials for each tail length. Flow speeds are indicated as cm·s^-1^ and BL· s^-1^ (body length). **(A)** ODBA variations in the myliobatid model. Models with tails maintained consistent ODBA across all speeds, without significant differences among both tail lengths (Posthoc T-test, p > 0.05), whereas models without tails exhibited significantly higher ODBA, particularly at speeds below 2.5 BL·s⁻¹ (25 cm·s⁻¹) (Posthoc T-test, p < 0.05). **(B)** Pitch range variations in the myliobatid model. Models without tails showed a greater pitch range than those with tails across all speeds (Posthoc T-test, p < 0.05). **(C)** Roll range variations in the myliobatid model. At speeds below 3 BL·s⁻¹ (30 cm·s⁻¹), models without tails exhibited a higher roll range than those with tails (Posthoc T-test, p < 0.05). **(D)** ODBA variations in the NACA 0012 model. NACA models without tails exhibited higher ODBA than those with tails at all speeds (Posthoc T-test, p < 0.05). **(E)** Pitch range variations in the NACA model. Models without tails significatively increased the pitch range at all speeds (Posthoc T-test, p < 0.05). **(F)** Roll range variations in the NACA model. Models without tails presented larger roll ranges at all speeds; being particularly high at lower speeds and decreasing as speed increased (Posthoc T-test, p < 0.05).

**Table 1.** Mean +/- standard error values for Pitch, Pitch range, Roll, Roll Range, and mean ODBA across speeds for the myliobatid model.

| Speed<br>cm/s | Speed<br>BL/s | Pitch Mean \| Pitch Range (°) |  |  |  |  |  | Roll Mean \| Roll Range (°) |  |  |  |  |  | ODBA (g) |  |  |
| --- | --- | --- | --- | --- | --- | --- | --- | --- | --- | --- | --- | --- | --- | --- | --- | --- |
|  |  | 30 cm |  | 15 cm |  | 0 cm |  | 30 cm |  | 15 cm |  | 0 cm |  | 30 cm | 15 cm | 0 cm |
| 10.8 | 1.1 | 27.2<br>±1.7 | 8.2<br>±1.4 | 24.9<br>±0.4 | 8.1<br>±1.2 | 17.2<br>±0.7 | 13.7<br>±0.4 | 2.1<br>±9.3 | 17.5<br>±3.6 | -8.5<br>±1.4 | 27.1<br>±3.5 | -4.1<br>±0.4 | 72.6<br>±0.4 | 0.02<br>±0.01 | 0.03<br>±0.006 | 0.48<br>±0.001 |
| 14.1 | 1.4 | 17.9<br>±4.1 | 8.7<br>±0.5 | 14.7<br>±1.0 | 5.6<br>±1.1 | 12.3<br>±0.6 | 17.7<br>±0.3 | -1.5<br>±1.8 | 9.3<br>±1.4 | -3.5<br>±0.8 | 10.6<br>±1.0 | -6.9<br>±1.5 | 52.4<br>±1.5 | 0.02<br>±0.002 | 0.03<br>±0.004 | 0.25<br>±0.02 |
| 17.3 | 1.7 | 10.3<br>±2.1 | 6.7<br>±0.5 | 8.8<br>±0.7 | 5.1<br>±0.3 | 8.1<br>±0.5 | 14.7<br>±0.5 | -3.2<br>±1.5 | 8.1<br>±0.8 | -3.5<br>±0.7 | 10.3<br>±1.0 | -5.7<br>±0.9 | 40.1<br>±0.9 | 0.02<br>±0.002 | 0.03<br>±0.002 | 0.16<br>±0.009 |
| 20.6 | 2.1 | 6.8<br>±1.2 | 6.2<br>±0.1 | 6.1<br>±0.7 | 5.1<br>±0.2 | 5.5<br>±0.4 | 10.3<br>±0.2 | -3.7<br>±1.6 | 8.7<br>±0.3 | -3.9<br>±0.9 | 8.6<br>±0.7 | -4.8<br>±1.7 | 28.1<br>±1.7 | 0.03<br>±0.0004 | 0.03<br>±0.001 | 0.12<br>±0.01 |
| 24.0 | 2.4 | 4.8<br>±1.0 | 6.7<br>±0.3 | 4.3<br>±0.7 | 6.8<br>±0.1 | 4.2<br>±0.4 | 10.4<br>±1.4 | -4.9<br>±1.6 | 9.1<br>±0.7 | -5.3<br>±1.1 | 9.6<br>±0.6 | -5.5<br>±1.8 | 18.5<br>±1.8 | 0.03<br>±0.0009 | 0.03<br>±0.001 | 0.06<br>±0.005 |
| 27.2 | 2.7 | 3.9<br>±0.9 | 8.9<br>±0.7 | 3.5<br>±0.6 | 9.5<br>±0.4 | 3.3<br>±0.4 | 13.1<br>±1.1 | -8.7<br>±1.2 | 10.3<br>±0.6 | -8.8<br>±0.8 | 10.9<br>±0.5 | -7.7<br>±1.8 | 17.7<br>±1.8 | 0.04<br>±0.002 | 0.04<br>±0.001 | 0.08<br>±0.01 |
| 30.5 | 3.1 | 3.3<br>±0.7 | 9.6<br>±0.5 | 2.9<br>±0.6 | 11.6<br>±0.8 | 2.8<br>±0.4 | 14.6<br>±0.7 | -11.7<br>±1.4 | 11.3<br>±1.1 | -11.6<br>±1.2 | 10.5<br>±0.4 | -10.0<br>±1.5 | 14.8<br>±1.5 | 0.04<br>±0.001 | 0.05<br>±0.001 | 0.07<br>±0.005 |
| 33.8 | 3.4 | 3.2<br>±0.7 | 10.7<br>±0.6 | 2.8<br>±0.6 | 10.9<br>±0.6 | 2.5<br>±0.4 | 13.1<br>±0.8 | -14.9<br>±1.6 | 11.2<br>±0.4 | -14.4<br>±1.4 | 10.2<br>±0.5 | -12.2<br>±0.9 | 11.3<br>±0.9 | 0.05<br>±0.002 | 0.04<br>±0.0001 | 0.06<br>±0.01 |
| 37.0 | 3.7 | 3.5<br>±0.6 | 10.4<br>±1.0 | 3.1<br>±0.7 | 11.9<br>±0.7 | 2.6<br>±0.4 | 13.3<br>±0.7 | -18.9<br>±1.9 | 9.8<br>±0.8 | -18.3<br>±1.8 | 9.6<br>±0.4 | -14.8<br>±0.6 | 10.5<br>±0.6 | 0.04<br>±0.002 | 0.04<br>±0.001 | 0.06<br>±0.005 |
| 40.3 | 4.0 | 4.3<br>±0.9 | 9.7<br>±0.4 | 3.9<br>±0.9 | 9.8<br>±0.2 | 2.6<br>±0.4 | 12.8<br>±0.7 | -24.0<br>±2.6 | 10.4<br>±0.9 | -23.1<br>±2.4 | 9.4<br>±1.4 | -18.5<br>±0.2 | 7.9<br>±0.2 | 0.04<br>±0.002 | 0.04<br>±0.0001 | 0.05<br>±0.002 |
| 43.6 | 4.3 | 5.6<br>±1.2 | 9.2<br>±0.1 | 5.2<br>±1.1 | 8.8<br>±0.2 | 2.9<br>±0.5 | 11.1<br>±0.7 | -30.2<br>±3.3 | 10.9<br>±1.5 | -29.5<br>±2.7 | 9.4<br>±0.4 | -23.3<br>±0.2 | 9.3<br>±0.2 | 0.03<br>±0.001 | 0.03<br>±0.001 | 0.04<br>±0.002 |
| 46.9 | 4.6 | 8.2<br>±2.9 | 8.8<br>±0.3 | 6.6<br>±1.5 | 9.0<br>±0.2 | 3.9<br>±0.6 | 9.6<br>±0.3 | -39.9<br>±7.6 | 12.7<br>±2.5 | -35.6<br>±3.8 | 10.1<br>±0.4 | -28.1<br>±0.1 | 9.3<br>±0.1 | 0.03<br>±0.0009 | 0.03<br>±0.002 | 0.04<br>±0.001 |

Tail length (0, 15, 30cm) had only minor effects on the posture of the models (pitch, roll) during testing (Fig. 6). In general, tail length did not significantly affect either model’s roll (myliobatid: F_2,134_ = 0.966, p = 0.38; NACA: F_2,132_ = 1.02, p = 0.36). For the NACA model, tail length did not significantly affect average pitch (F_2,132_ = 1.28, p=0.28) (Fig. 5, Table S1), regardless of speed (F_32,132_ = 0.13, p >0.99) (Fig 6B). In contrast, for the myliobatid model, tail length significantly impacted body pitch (F_2,134_=4.9, p = 0.009), although this effect was not uniform across speeds (F_28,134_ = 2.12, p = 0.002) (Fig. 6A). At the lowest speeds (1 BL ·s^-1^), myliobatid models with tails had higher mean pitch values (30 cm tail = 27.20 ± 1.7; 15 cm tail = 24.93 ± 0.4) than the model without a tail (0 cm tail = 17.25 ± 0.7) (Tukey HSD, 0:15cm: p < 0.001; 0:30cm: p = 0.005) (Table 1; Fig. 6A). However, as flow speed increased, no significant differences in body pitch were observed between myliobatid models with different tail lengths (Tukey HSD, p > 0.1) (Fig. 6A).

There were differences in the average posture of the NACA and myliobatid models. The myliobatid model displayed higher pitch values than the NACA model (0cm: F_1,78_ =261, p <0.001; 15cm: F_1,78_ =172, p <0.001; 30cm: F_1,78_ =119, p <0.001), and this was true at all speeds (10-50 cm·s^-1^, pairwise T-Test p < 0.05) (Table 1, Table S1). As the myliobatid models rolled onto their sides at increasing speed (i.e. tilted to the right) and the NACA models did not, there were significant differences in the roll posture between the models (0cm: F_1,78_ =3329, p <0.001; 15cm: F_1,78_ =279, p <0.001; 30cm: F_1,78_ =148, p <0.001). However, at low speeds (<225 rpm), there were no differences between the average roll position of the NACA and myliobatid models (Paired T-test, p>0.05).

### Model stability

To quantify model stability, we used three variables that described the movement of the model: 1) the overall dynamic body acceleration (ODBA) combines the dynamic acceleration of an animal’s body across X, Y, and Z axis; 2) Pitch range, that quantify the movement of the model in Y-Z (anterior-posterior); and 3) Roll range, that quantify the movement of the model in X-Y (medial-lateral). Flow speed and tail length had a large impact on all three variables, and therefore, an effect on the model stability (Fig. 7; Tables S2–S4).

Regardless of flow speed, both the myliobatid and NACA without tails (0 cm) moved significantly more (higher ODBA) than either of the models with tails (15 & 30cm; Posthoc T-test p<0.05) (Fig. 7A, D; Table 1; Table S2–S4). These stability differences between tailed and tailless models were most pronounced at the lowest flow speed (1.0 BL·s^-1^) in which the tailless models had an order of magnitude higher ODBA (myliobatid – 0 cm tail = 0.48 ± 0.001g; NACA model – 0 cm = 0.28± 0.008g) than the tailed models (ODBA < 0.03g) (Table 1, Table S1). As flow speed increased, the ODBA of the tailless NACA and myliobatid models decreased (indicating decreased movement) (Fig. 7A,D). However, as the flow speed increased (speed > 3 BL ·s^-1^), the myliobatid model approached ODBA values similar to myliobatid models with tails (Fig. 7A; Table 1), the tailless NACA model maintained a higher ODBA than those with tails for all speeds (Fig. 7D; Table S1).

Doubling the tail length from 15 to 30cm, did not significantly change the ODBA of either the NACA or myliobatid models in over 90% of tested speeds (Posthoc T-Test p>0.05) (Tables S2– S4). Additionally, while there were significant differences between the myliobatid and NACA model’s ODBA (0cm: F_1,78_ =53, p <0.001; 15cm: F_1,78_ =23, p <0.001; 30cm: F_1,78_ =24, p <0.001), at speeds below 2.4 BL·s^-1^ the NACA and myliobatid models with 15 and 30cm tails did not differ from each other (Posthoc T-Test: p>0.05).

To quantify how the models were moving, we further assessed the variability in model posture at each speed using pitch range and roll range variables. Similar to ODBA, models with tails (15 and 30 cm) displayed similar anterior-posterior (pitch range; Fig. 7B, E) and medial-lateral (roll range; Fig. 7C, F) postural stability to each other, and significant differences (Posthoc T-test, p<0.05) between tail lengths (15 and 30cm) were only observed in 3 out of 56 posthoc pairwise comparisons. Furthermore, while there was significant variation in pitch range and roll range stability with speed (Table S2–S4), for the tailed models there were not clear trends displaying a similar (< 4° difference) pitch range and roll range across speeds (Fig. 7B–F; Table 1, Table S1).

In contrast, for both myliobatid and NACA, tailless models differ significantly from both tailed modes, presenting a higher pitch and roll range (Posthoc T-Test: p < 0.05). For pitch range (anterior-posterior stability), this result is evidenced by a significant larger pitch range (myliobatid – 0 cm tail = 13.7 ± 0.4°, NACA – 0 cm tails = 17.6 ± 1.1°) compared to the tailed models (myliobatid – 15, 30 cm tail ≈ 8°, NACA – 15, 30 cm tails < 7°; Table 1, Table S1). However, these differences were more pronounced at lower speeds (speed < 2.06 BL· s^-1^). Roll range (medial-lateral stability) was the variable most affected by the tail length, especially at lower speeds where the roll range of models without tail were ±25° (myliobatid – 0 cm tail = 27.08 ± 3.5°, NACA – 0 cm tails = 65.78 ± 8.7°; Table 1, Table S1), while the tailed models’ oscillations were < ± 5.5° (Fig. 7C, F; Table 1, Table S1). As flow speed increased the roll range of the tailless models was reduced (and, therefore, the medial-lateral stability increased) and the tailless myliobatid model’s roll range was not significantly different than the tailed models once flow speed exceeded 2.4 BL· s^-1^. However, the tailless NACA model maintained an elevated roll range compared to the tailed models until flow speeds exceeded 4.0 BL· s^-1^. Furthermore, while the roll stability of the tailless myliobatid model increased rapidly with flow speed (Fig. 7C), the tailless NACA model’s stability increased more linearly, with the NACA model having a significantly higher roll range at flow speeds between 1.7 BL· s^-1^ and 3.7 BL· s^-1^ (Fig. 7F)

We also investigated the minimum tail length required to stabilize the models (Fig. 8). For the myliobatid model, tail lengths ranging from 3 times to 0.9 times the body length (STL 3 – 0.9 BL) effectively stabilized the model, with no observed differences among these lengths (Fig. 8A). However, when the STL was shorter than 0.9 BL, the myliobatid model began to destabilize critically, reflected in a rapid increase of ODBA with decreasing tail length (Fig. 8A). A similar trend was observed for the NACA model, although the increase in ODBA values was more gradual as the tail shortened (Fig. 8B). For the NACA model, the minimum STL to stabilize the model was 1.2 (12 cm tail length). Beyond this length, ODBA values progressively increased, peaking in the absence of a tail (Fig. 8B).

**Figure 8.**
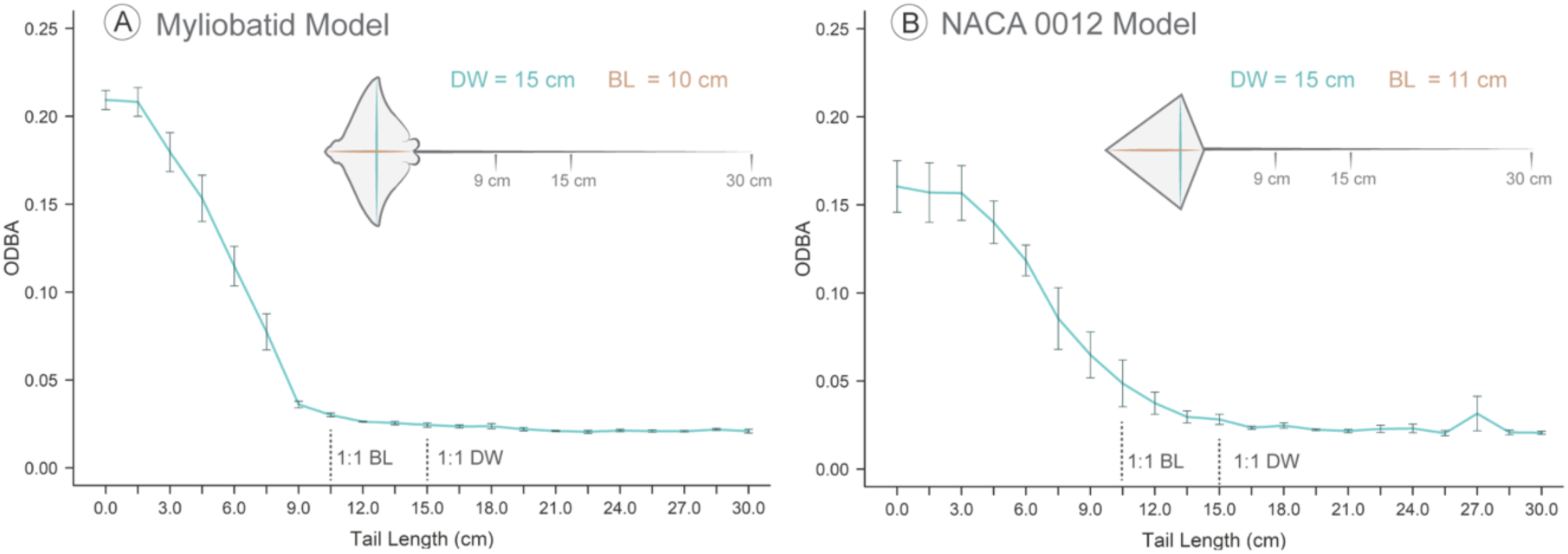
ODBA variation with progressive reduction in tail length. **(A)** ODBA values for the myliobatid model as the tail was progressively shortened by 1.5 cm. ODBA remained consistently below 0.05g when the tail length was greater than 9 cm (0.9 BL). However, when the tail was shorter than 0.9BL, ODBA drastically increased as shorter was the tail, reaching maximum values at 2.7 BL. **(B)** For the NACA 0012 model, ODBA remained relatively stable until the tail length dropped below 11 cm (1 BL), after which ODBA progressively increased, reaching the highest values with a tail length of 3 cm (0.27 BL). Note that BL differ for each model, being 10 cm for the Myliobatid model and 11 cm for the NACA 0012 model. Error bars represent standard errors from the three trials conducted for each model. All experiments were performed at a flow speed of 1.4 BL·s⁻¹ (14 cm s⁻¹). BL = Body length, DW = Disc width

## DISCUSSION

In this study, we examine variation in structure and function of the long, slender tails across the families of the order Myliobatiformes which use oscillatory pectoral fin locomotion, and explore how these tails provide passive stability in models with the large pectoral fins fixed in an extended position. Our results clearly indicate that models without tails are significantly less stable than models with tails, exhibiting greater roll (medial-lateral motion), pitch (anterior-posterior motion), and overall dynamic body acceleration (ODBA). Additionally, we identified that the minimum tail length required to stabilize the Myliobatid-based model was 0.9 times the body length. Tails exceeding this threshold provided no additional stabilization, as models with tail lengths up to 3 times the body length or 2 times the disc width—the maximum tested in this study—displayed similar levels of stability (Fig. 8). Interestingly, these findings align with the variation in relative tail length observed across myliobatid species (Fig. 3). The minimum tail length of 0.9 times the body length (BL) is characteristic of the giant manta ray (*Mobula birostris*), whereas the maximum relative tail length of 4.6 times the body length is found in the spotted eagle ray (*Aetobatus narinari*). Notably, no myliobatid species examined had a tail length shorter than 0.9 BL. These differences in tail length reflect the dual role of tails in myliobatids: our results suggest that the tail is crucial for passively stabilizing the body during gliding locomotion when the pectoral fins are not moving. Our finding that longer tails do not provide additional stabilization suggest their involvement in other functions beyond stability, such as sensory or defensive roles.

### Tails as a body stabilizing structures

The distinctive body morphology of myliobatids, with their broad, tapered pectoral fins, generates thrust through lift-based mechanisms and achieves a high lift-to-drag ratio, enabling effective propulsion during the fins’ stroke cycles (Fish et al., 2016; Heine, 1992; Liu et al., 2015; Russo et al., 2015). However, the function during locomotion of the whip-like tails, a distinctive feature of myliobatids, has often been overlooked.

Unlike other marine vertebrates, such as teleost fishes and sharks, myliobatids do not use their tails for propulsion. Instead, their tails are relatively rigid, as they are supported by a long, mineralized rod of fused vertebrae (the caudal synarcual) that minimizes bending (Chaumel and Lauder, 2025). Additionally, myliobatids maintain the tail passively stiffened and straight, as musculature is limited to the attachment point with the body, leaving the remainder of the tail almost devoid of muscles (Chaumel and Lauder, 2025) (Movies S1–S2). To our knowledge, this internal tail anatomy is distinctive of myliobatids and it has not been observed in any other marine or terrestrial vertebrate, including the related species that also are characterized by a whip-tail. Unlike myliobatids, other families within the order Myliobatiformes are characterized by benthic ecologies and undulatory locomotion. Some species from these families, especially Dasyatidae (dasyatids), Potamotrygonidae (freshwater stingrays) and Gymnuridae (butterfly rays), are also characterized by whip-like tails. However, although relatively unknown, their tails are characterized by a segmented vertebral column instead of a caudal synarcual and, in several species, the tails can be also associated with fins (Fontanella et al., 2013; Last et al., 2016; Nishida, 1990; Rosenberger, 2001; Rosenberger and Westneat, 1999). Having segmented vertebrae would allow for a higher degree of motion, more suited to other functions related to the lifestyle of these species, including defense (by supporting and moving the barb) or counterbalancing during undulatory locomotion near the sea bottom. However, further research is needed to assess the extent to which a segmented vertebral column within the tail relates to locomotor ecology.

The function of a tail in mediating body stability can be seen in analyses of flight by kites (Borobia-Moreno et al., 2021; Marvin, 1897; NASA, 2021). In kites, long and slender tails play a crucial aerodynamic role by passively increasing the stability of the kites while flying by reducing roll, yaw and pitch (Marvin, 1897; NASA, 2021). This resembles the effect of tail in our myliobatid and NACA models, where tails significatively increased stability (ODBA) by reducing anterior-posterior (pitch) and medial-lateral movements (roll). As a result, we hypothesize that the same physical phenomena that apply to the tails in kites applies to myliobatids during gliding locomotion, an effect that we term the *kite hypothesis*. The physical phenomena generated by long and slender tails in kites are based on adding drag, which produces a restoring torque and, ultimately, stabilizes the center of pressure (point where forces are balanced).

Tails increase body stability (Fig. 9). Without tails, the center of pressure of our models has a larger range of anterior-posterior movement (which increases pitch range) and medial-lateral movements (which increases roll range). In contrast, long and slender tails add drag posterior to both the center of mass and pressure, a phenomenon known as drag-based stability (Marvin, 1897; NASA, 2021; Thomas and Taylor, 2001). By adding drag, tails act to damp oscillations, generating a restoring torque (rotational force) which reduces the range of movement of the center of pressure and maintains it more centered in the body (Fig. 9). Reducing the range of movement of the center of pressure reduces the response of the body to disturbances in roll and pitch. As the tail increases in length, it produces a more effective torque in restoring balance due to increased frictional drag force: total friction drag is a function of tail length and hence surface area. This effect was observed in our study, as tails lengths ranging from 0.9 to 3 times the body length (maximum length tested in our experiments) equally stabilized the models, with no significant difference in their effects on pitch and roll range, or ODBA. However, when the tail length dropped below 0.9 times body length, destabilization occurred, increasing pitch and roll range and ODBA. This suggests that a tail shorter than this threshold does not experience sufficient drag and is therefore unable to generate sufficient restoring torque to stabilize the body, preventing it from damping oscillations and, therefore, not returning the animal to its original position (Fig. 9).

**Figure 9.**
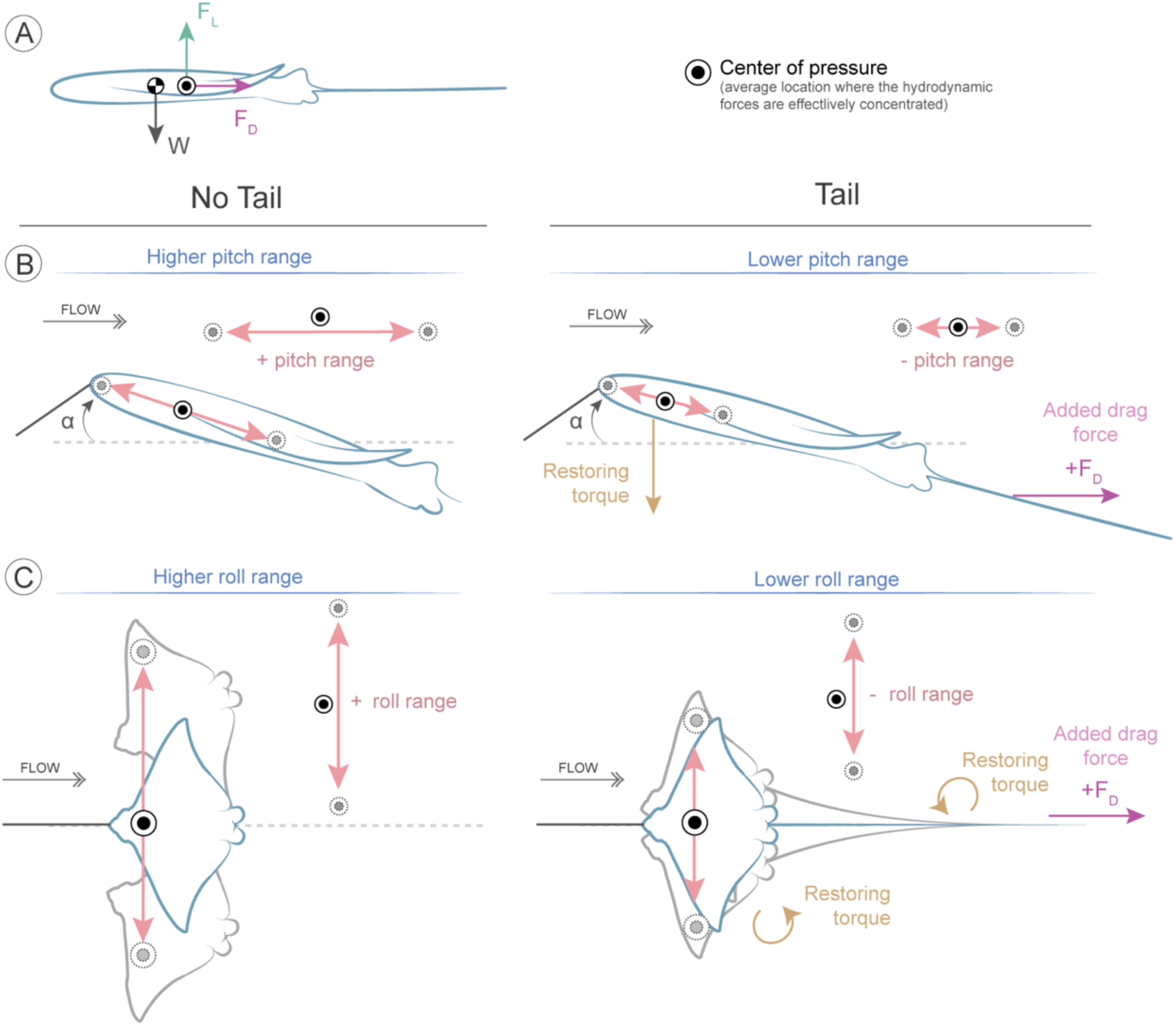
Schematic interpretation of the body stabilizing effect of the whip-like tail of myliobatids, illustrating the kite hypothesis of tail stability. **(A)** Simplified representation of the forces acting on a body gliding through flow with the pectoral fins extended. Forces include lift force (F_L_) and drag force (F_D_), which balance at the center of pressure 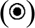, and weight (W), which is balanced at the center of mass 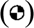. **(B)** Effects of the tail on pitch stability. We hypothesize that the tail helps stabilize the center of gravity in anterior-posterior movement (pitch range). In the absence of a tail, the model exhibits a higher pitch range. However, a long, slender tail reduces pitch fluctuations by generating drag at the rear, damping forces, and acting as a restoring torque, ultimately leading to a more stable posture. Since the tail is lightweight, it does not significantly alter the mass distribution or angle of attack but experiences friction drag as a function of its surface area. **(C)** Effect of the tail on roll stability. The tail reduces roll range by limiting lateral-medial (roll range) movement around the center of gravity. Models without tails experience greater roll fluctuations, whereas those with tails generate additional drag and damping forces, acting as a restoring torque that minimizes lateral-medial oscillations. To effectively reduce both pitch and roll range, the tail must be at least 0.9 BL in length.

In contrast, tails have minor effects on model posture. The tails of myliobatids, as well as the tails used in our study, are relatively light compared to the body of the animal (1.3% in real myliobatids and 2.3% in our models). Therefore, longer tails do not add significant weight that could affect the position of the center of gravity (center of mass), which is located anterior to the fin tips (Fontanella et al., 2013). This is evident in our results, as models with larger tails displayed only minor differences in body posture and only at the slowest speeds. Lastly, myliobatid tails are relatively rigid, which limits their bending as rays swim and glide (Chaumel and Lauder, 2025) (Movies S1– S2).

Finally, we emphasize that while our experimental analysis of the effect of tail length on body stability is most relevant to gliding behavior, similar effects could be seen during flapping propulsion. Animals often use appendage movements to stabilize the head and visual field even as thrust forces are generated by limbs or fins (Dunbar, 2004; Fish et al., 2003; Kress and Egelhaaf, 2012; Lapsansky and Tobalske, 2019; Pozzo et al., 1990; Xiong and Lauder, 2014). A long and relatively stiff tail held behind the body could act to dampen vertical center of mass oscillations during oscillatory propulsion generated by either flapping or undulating pectoral fins in rays. The role of the tail in maintaining body stability during active propulsion is an important area for future study.

### Tail effect on body morphology

The effects of body morphology were tested by comparing a model based on a myliobatid body to a model based on a classic NACA 0012 airfoil body. Both tail lengths (15 and 30 cm) equally stabilized both models, resulting in similar ODBA values across all flow speeds. This indicates that light, long, and slender tails are crucial structures to and function to increase stability similarly in different body shapes. However, without tails, models behave differently, which demonstrates the effect of body morphology. Without tails, the NACA model showed lower ODBA values than the myliobatid model, indicating that the NACA shape is more stable than a myliobatid triangular shape. However, flow speed had different impact on stabilizing both models without tails. While the tailless myliobatid model stabilized at flow speeds higher than ∼2.4 BL·s^-1^, showing similar ODBA values than models with tails, the tailless NACA model always maintained higher ODBA values than models with tails across all speeds. This indicates that the myliobatid morphology is able to stabilize itself with high flow speeds, while the NACA model cannot. In addition, the minimum tail length required to stabilize the models differed between NACA and myliobatid body models. The minimum tail length to stabilize the myliobatid model was 0.9 body lengths; at tail lengths less than this value, the model became unstable. For NACA body model, instability began at tail lengths less than 1.2 times body length (Fig. 8).

### The role of the tail length in myliobatid biology

In this study, we found interesting correspondences between our experimental results on body models and measurements of tail length in myliobatid rays. *Mobula birostris* presented the shortest tails relative to the body, with a minimum value measured of 0.9 times body length. Similarly, 0.9 times body length was the minimum tail length value required to stabilize experimentally the myliobatid model (Fig. 8). The fact that none of the myliobatid species measured had tails shorter than this value suggests that tails play an important role in maintaining body stability during the gliding phase (non-active locomotion) of myliobatid rays. Our experimental tests also show that tails greatly increase body stability at speeds between 1 and 2.5 BL·s^-1^. This result agrees with the speed range recorded for swimming and gliding myliobatids, as reported by (Fish et al., 2016; Fong et al., 2022; Fontes et al., 2018; Fontes et al., 2022) who noted a speed range of 0.2 to 1.25 BL·s^-1^ for *M. birostris*.

In contrast, tails longer than 0.9 BL did not provide additional stability, suggesting that in species with tails exceeding this threshold, tails may be involved in additional functions. This idea is supported by the observed variation in relative tail length across myliobatid species and throughout ontogeny (Fig. 3). Although myliobatids are described as “pelagic rays”, migratory species able to efficiently swim long distances and over long periods of time, species differ considerably in ecology and lifestyle. Species from the families *Aetobatidae* (e.g., the spotted eagle ray *Aetobatus narinari*), *Rhinopteridae* (e.g., the cownose ray *Rhinoptera bonasus*), and *Myliobatidae* (bat eagle ray *Myliobatis californica*) are benthopelagic predators, since they mainly predate on mollusks buried in the sand (Cahill et al., 2023; Collins, 2005; DeGroot et al., 2021; Fisher, 2010; Sasko et al., 2006; Schluessel et al., 2010). In contrast, devil and manta rays (family *Mobulidae*) are filter-feeders that they feed on plankton that live suspended in the water column (and are specialized to detect and predate on them using their cephalic lobes) and, therefore, are not associated with the sea-bottom (Beale et al., 2019; Couturier et al., 2012; Sampson et al., 2010). However, although devil and manta rays present similar feeding behavior, they differ in size; while manta rays (*M. birostris* and *Mobula alfredi*) are some of the largest fishes on the ocean, devil rays are similar in size to other myliobatids (Last et al., 2016).

Our results indicate that tail length correlates with body size, regardless of feeding ecology (Fig. 3). While larger specimens have the shortest tails, smaller species, such as cownose, eagles, and devil rays, have similar relative tail lengths. These findings suggest a possible correlation between tail length and the capacity to maneuver (defined as the ability to change position, direction, and orientation while maintaining stability). Smaller rays are more agile and capable of rapid turns compared to manta rays, and relatively larger tails may help in maintaining stability during rapid turns. Additionally, myliobatid tails may be related to predator avoidance. Smaller rays are also more vulnerable to predators than larger rays (e.g., sand sharks, hammerheads and killer whales are known predators of several myliobatid species) (Couturier et al., 2012; Flowers et al., 2021; Higuera-Rivas et al., 2023). The mechanosensory system of the tail – which contains an extensive lateral line (Chaumel and Lauder, 2025) – may play a crucial role in improving predator detection: longer tails extend the spatial range for detecting predators approaching from behind, giving rays sufficient time to scape. Previous work has shown that rays initiate an escape response when a sudden water movement is produced near the tail (Chaumel and Lauder, 2025). Finally, the tail may play a role in inter-individual communication and may facilitate social interactions during the formation of large schools (common in myliobatid species) (Grancagnolo and Arculeo, 2021; Palacios et al., 2024; Rogers et al., 1990) and mating, as suggested by observations of myliobatids in captivity in which an individual will follow a leader while maintaining physical contact the tail leaders tail (Movie S2).

## Supporting information

Movie S1. Aetobatus narinari (spotted eagle ray) gliding at the Georgia Aquarium.

Movie S2. Mobula birostris (giant manta ray) gliding at the Georgia Aquarium.

## LIST OF SYMBOLS AND ABBREVIATIONS USED

AR: Aspect Ratio
BL: Body Length
DW: Disc Width
ODBA: Overall Dynamic Body Acceleration
STL: Standard Tail Length
TL: Tail Length

## ACKNOWLEDGEMENTS

We sincerely thank Meaghan Sorce and Andrew Williston from the Ichthyology Collection of the Museum of Comparative Zoology at Harvard University, Diego Vaz from the Natural History Museum in London, Kevin W. Conway and Sabri Amrani-Khaldi from Texas A&M University, and Will White from CSIRO in Hobart for their invaluable assistance measuring, taking and sharing images of the specimens from the collections. We especially thank Yangfan Zhang for insightful discussions and for suggesting the kite hypothesis, which greatly contributed to understanding the results of this study.

## DATA AND RESOURCE AVAILABILITY

Biologger raw data and specimens analyzed ID are publicly available at Mendeley Repository.

