## Supplementary material for "The tail of myliobatid rays controls body stability": Sup. Materials

**Table S1**. Mean +/- standard error values of Pitch, Pitch range, Roll, Roll Range, and mean ODBA across speeds for the NACA model.

| Speed cm/s | Speed BL/s | Pitch Mean \| Pitch Range (º) | | | | | | Roll Mean \| Roll Range (º) | | | | | | ODBA (g) | | |
|---|---|---|---|---|---|---|---|---|---|---|---|---|---|---|---|---|
| | | 30 cm | | 15 cm | | 0 cm | | 30 cm | | 15 cm | | 0 cm | | 30 cm | 15 cm | 0 cm |
| 10.8 | 1.1 | 13.1 ±1.1 | 4.9 ±0.9 | 13.0± 1.5 | 7.0 ±0.4 | 12.3± 0.7 | 17.2 ±1.1 | 1.9 ±1.2 | 5.3 ±0.7 | 0.6 ±0. 7 | 7.5 ±1.6 | 3.2 ±8.7 | 65.7 ±8.7 | 0.01 ±0.000 1 | 0.02 ±0.006 | 0.27 ±0.008 |
| 14.1 | 1.4 | 6.8 ±0.9 | 4.8 ±0.6 | 6.4 ±1.0 | 6.1 ±0.5 | 6.1 ±0.6 | 19.8 ±1.9 | 1.5 ±1.5 | 5.6 ±0.8 | 0.5 ±0.8 | 10.7 ±0.8 | 1.1 ±6.8 | 63.0 ±6.8 | 0.01 ±0.001 | 0.04 ±0.004 | 0.21 ±0.01 |
| 17.3 | 1.7 | 3.5 ±0.7 | 4.7 ±0.5 | 3.4 ±0.7 | 4.9 ±0.2 | 3.2 ±0.5 | 21.9 ±0.9 | 1.0 ±1.6 | 7.1 ±0.9 | 0.2 ±0.9 | 9.1 ±0.7 | 5.7 ±5.3 | 60.6 ±5.3 | 0.02 ±0.001 | 0.02 ±0.001 | 0.18 ±0.003 |
| 20.6 | 2.1 | 1.8 ±0.6 | 5.0 ±0.1 | 1.8 ±0.7 | 5.0 ±0.1 | 1.5 ±0.4 | 19.4 ±1.9 | 0.9 ±1.7 | 8.2 ±1.0 | 0.2 ±1.0 | 9.1 ±0.9 | -1.4 ±6.1 | 57.2 ±6.1 | 0.02 ±0.000 1 | 0.02 ±0.001 | 0.15 ±0.07 |
| 24.0 | 2.4 | 0.9 ±0.6 | 5.3 ±0.2 | 0.9 ±0.7 | 5.5 ±0.2 | 0.5 ±0.4 | 17.0 ±1.3 | 1.1 ±1.8 | 7.6 ±1.3 | 0.5 ±1.3 | 8.7 ±0.7 | 5.7 ±3.1 | 48.5 ±3.1 | 0.02 ±0.000 2 | 0.02 ±0.001 | 0.14 ±0.003 |
| 27.2 | 2.7 | 0.2 ±0.7 | 5.8 ±0.2 | 0.2 ±0.7 | 5.3 ±0.4 | -0.1 ±0.4 | 16.3 ±0.2 | 1.6 ±1.9 | 8.8 ±1.5 | 0.9 ±1.5 | 9.2 ±1.0 | 1.5 ±5.0 | 52.0 ±5.0 | 0.03 ±0.001 | 0.02 ±0.001 | 0.13 ±0.07 |
| 30.5 | 3.1 | -0.2 ±0.7 | 5.7 ±0.3 | -0.2 ±0.8 | 5.8 ±0.2 | -0.5 ±0.4 | 17.5 ±1.3 | 2.0 ±2.1 | 8.5 ±1.7 | 1.7 ±1.7 | 8.8 ±0.9 | 1.3 ±5.9 | 46.0 ±5.9 | 0.03 ±0.002 | 0.03 ±0.001 | 0.12 ±0.003 |
| 33.8 | 3.4 | -0.6 ±0.8 | 5.6 ±0.1 | -0.6 ±0.8 | 5.8 ±0.2 | -0.9 ±0.4 | 15.7 ±0.5 | 2.4 ±2.5 | 7.9 ±1.9 | 2.1 ±1.9 | 9.1 ±1.0 | -1.6 ±5.6 | 47.5 ±5.6 | 0.03 ±0.002 | 0.03 ±0.000 2 | 0.12 ±0.03 |
| 37.0 | 3.7 | -0.8 ±0.8 | 6.1 ±0.2 | -0.9 ±0.8 | 6.2 ±0.1 | -1.2 ±0.4 | 13.3 ±1.2 | 2.5 ±3.5 | 8.4 ±2.3 | 2.3 ±2.3 | 10.4 ±1.4 | 2.5 ±6.8 | 39.8 ±6.8 | 0.03 ±0.002 | 0.03 ±0.002 | 0.11 ±0.006 |
| 40.3 | 4.0 | -0.7 ±1.2 | 5.6 ±0.3 | -1.1 ±0.9 | 5.7 ±0.2 | -1.4 ±0.4 | 14.2 ±1.7 | 3.3 ±4.2 | 8.7 ±2.6 | 2.5 ±2.6 | 9.0 ±1.3 | -2.7 ±8.0 | 31.5 ±8.0 | 0.03 ±0.002 | 0.03 ±0.000 2 | 0.11 ±0.01 |
| 43.6 | 4.3 | -0.7 ±1.4 | 5.6 ±0.1 | -0.9 ±1.1 | 6.4 ±0.6 | -1.6 ±0.5 | 13.1 ±1.9 | 3.6 ±4.8 | 8.4 ±3.7 | 3.3 ±3.7 | 9.7 ±0.2 | -1.6 ±8.7 | 29.4 ±8.7 | 0.03 ±0.001 | 0.03 ±0.002 | 0.10 ±0.002 |
| 46.9 | 4.6 | -0.7 ±1.5 | 6.0 ±0.1 | -0.8 ±1.3 | 6.2 ±0.4 | -1.7 ±0.6 | 11.5 ±1.4 | 3.6 ±5.3 | 8.9 ±4.5 | 3.8 ±4.5 | 9.1 ±0.8 | 2.5 ±6.7 | 23.3 ±6.7 | 0.03 ±0.000 1 | 0.03 ±0.002 | 0.08 ±0.001 |

**Table S2**. Results of one-way ANOVA on the myliobatid and NACA model.

| Model | Variable | Factor | Df | Sum sq | Mean sq | F-value | p-value |
|---|---|---|---|---|---|---|---|
| Myliobatid | Pitch Range | Tail length | 2 | 0.38 | 0.19 | 5.77 | 0.0396* |
| | | Flow speed | 15 | 7.12 | 0.47 | 14.08 | < 0.001*** |
| | | Tail length:Flow speed | 28 | 4.09 | 0.15 | 4.33 | < 0.001*** |
| | | Residuals | 134 | 4.52 | 0.03 | | |
| | Roll Range | Tail length | 2 | 3.29 | 1.64 | 3.44 | 0.0349* |
| | | Flow speed | 15 | 33.52 | 2.23 | 4.67 | < 0.001*** |
| | | Tail length:Flow speed | 28 | 28.14 | 1.00 | 2.10 | 0.0027 ** |
| | | Residuals | 134 | 64.06 | 0.47 | | |
| | ODBA | Tail length | 2 | 0.47 | 0.24 | 539.43 | < 0.001*** |
| | | Flow speed | 15 | 0.44 | 0.03 | 66.94 | < 0.001*** |
| | | Tail length:Flow speed | 28 | 1.12 | 0.04 | 90.95 | < 0.001*** |
| | | Residuals | 134 | 0.06 | 0.001 | | |
| NACA | Pitch Range | Tail length | 2 | 1.29 | 0.65 | 588.73 | < 0.001*** |
| | | Flow speed | 15 | 0.21 | 0.01 | 12.78 | < 0.001*** |
| | | Tail length:Flow speed | 30 | 0.36 | 0.01 | 12.78 | < 0.001*** |
| | | Residuals | 132 | 0.16 | 0.001 | | |
| | Roll Range | Tail length | 2 | 14.92 | 7.46 | 392.63 | < 0.001*** |
| | | Flow speed | 15 | 1.82 | 0.12 | 6.38 | < 0.001*** |
| | | Tail length:Flow speed | 30 | 3.24 | 0.11 | 5.69 | < 0.001*** |
| | | Residuals | 132 | 2.51 | 0.02 | | |
| | ODBA | Tail length | 2 | 0.54 | 0.27 | 853.63 | < 0.001*** |
| | | Flow speed | 15 | 0.11 | 0.01 | 23.95 | < 0.001*** |
| | | Tail length:Flow speed | 30 | 0.17 | 0.005 | 18.16 | < 0.001*** |
| | | Residuals | 132 | 0.05 | 0.0003 | | |

**Table S3.** T-test post-hoc results for the one-way ANOVA on the myliobatid model. Colored squares indicate statistical significance.

| | Pitch range | | | Roll Range | | | ODBA | | |
|---|---|---|---|---|---|---|---|---|---|
| Speed (rpm) | 0 – 15 | 0 – 30 | 15 – 30 | 0 – 15 | 0 – 30 | 15 – 30 | 0 – 15 | 0 – 30 | 15 – 30 |
| 100 | 0.016* | 0.025* | 0.988 | < 0.001* | < 0.001* | 0.111 | < 0.001* | < 0.001* | 0.824 |
| 125 | < 0.001* | < 0.001* | 0.064 | 2.099 | < 0.001* | 0.487 | 0.001* | 0.001* | 0.131 |
| 150 | < 0.001* | < 0.001* | 0.051 | 6.790 | < 0.001* | 0.152 | < 0.001* | < 0.001* | 0.380 |
| 175 | < 0.001* | < 0.001* | 0.007* | < 0.001* | 0.001* | 0.922 | 0.003* | 0.0035* | 0.192 |
| 200 | 0.081 | 0.077 | 0.902 | 0.013* | 0.010* | 0.619 | 0.008* | 0.007* | 0.142 |
| 225 | 0.045* | 0.028* | 0.556 | 0.029* | 0.021* | 0.451 | 0.083 | 0.058 | 0.111 |
| 250 | 0.036* | 0.003* | 0.096 | 0.065 | 0.120 | 0.574 | 0.032* | 0.015* | 0.026* |
| 275 | 0.098 | 0.077 | 0.824 | 0.359 | 0.880 | 0.242 | 0.001* | 0.009* | 0.497 |
| 300 | 0.245 | 0.073 | 0.287 | 0.591 | 0.520 | 0.918 | 0.009* | 0.007* | 0.777 |
| 325 | 0.023* | 0.020* | 0.955 | 0.027* | 0.081 | 0.395 | 0.001* | < 0.001* | 0.987 |
| 350 | 0.004* | 0.010* | 0.317 | 0.377 | 0.376 | 0.314 | 0.005* | 0.005* | 0.919 |
| 375 | 0.462 | 0.317 | 0.962 | 0.198 | 0.278 | 0.517 | 0.050 | 0.021* | 0.541 |
| 400 | 0.421 | 0.503 | 0.972 | 0.367 | 0.383 | 0.577 | 0.440 | 0.013* | 0.224 |

**Table S4.** T-test post-hoc results for the one-way ANOVA on the NACA model. Colored squares indicate statistical significance.

| | Pitch range | | | Roll Range | | | ODBA | | |
|---|---|---|---|---|---|---|---|---|---|
| Speed (rpm) | 0 – 15 | 0 – 30 | 15 – 30 | 0 – 15 | 0 – 30 | 15 – 30 | 0 – 15 | 0 – 30 | 15 – 30 |
| 100 | < 0.001* | < 0.001* | 0.105 | 0.005* | 0.006* | 0.278 | < 0.001* | < 0.001* | 0.163 |
| 125 | 0.004* | 0.002* | 0.166 | 0.004* | 0.003* | 0.005* | < 0.001* | < 0.001* | 0.006* |
| 150 | < 0.001* | < 0.001* | 0.824 | 0.002* | 0.001* | 0.102 | < 0.001* | < 0.001* | 0.076 |
| 175 | 0.005* | 0.004* | 0.992 | 0.003* | 0.003* | 0.605 | < 0.001* | < 0.001* | 0.386 |
| 200 | 0.002* | 0.002* | 0.524 | < 0.001* | < 0.001* | 0.436 | < 0.001* | < 0.001* | 0.538 |
| 225 | < 0.001* | < 0.001* | 0.384 | 0.002* | 0.003* | 0.706 | < 0.001* | < 0.001* | 0.840 |
| 250 | 0.002* | 0.002* | 0.715 | 0.007* | 0.007* | 0.828 | < 0.001* | < 0.001* | 0.852 |
| 275 | < 0.001* | < 0.001* | 0.324 | 0.005* | 0.005* | 0.440 | 0.002* | 0.002* | 0.888 |
| 300 | 0.008* | 0.007* | 0.5814 | 0.020* | 0.017* | 0.337 | < 0.001* | < 0.001* | 0.963 |
| 325 | 0.014* | 0.013* | 0.945 | 0.066 | 0.066 | 0.881 | 0.003* | 0.003* | 0.849 |
| 350 | 0.036* | 0.031* | 0.313 | 0.110 | 0.095 | 0.118 | 0.008* | 0.008* | 0.899 |
| 375 | 0.027* | 0.028* | 0.724 | 0.125 | 0.122 | 0.867 | 0.066 | 0.068 | 0.721 |
| 400 | 0.015* | 0.025* | 0.223 | 0.216 | 0.222 | 0.880 | 0.070 | 0.079 | 0.322 |

**Movie S1.** *Aetobatus narinari* (spotted eagle ray) gliding at the Georgia Aquarium.

**Movie S2.** *Mobula birostris* (giant manta ray) gliding at the Georgia Aquarium.
